# Evaluation of Detection Methods for Wastewater Surveillance of Antimicrobial-Resistant Bacteria from Healthcare Facilities

**DOI:** 10.64898/2025.12.14.694051

**Authors:** Ean Warren, James VanDerslice, L. Scott Benson, William J. Brazelton, Windy Tanner, Amanda K. Lyons, Florence Whitehill, Angela Coulliette-Salmond, Sydney Fusco, Jennifer Weidhaas

## Abstract

Carbapenem resistance is an urgent public health threat. Wastewater surveillance could support antimicrobial resistance monitoring at long-term care facilities (LTCFs). Feasibility of wastewater sampling (via composite or passive sampler, and sewer biofilm swabs) for carbapenemase genes (*bla*_KPC_, *bla*_VIM_, *bla*_OXA-48-like_, *bla*_NDM_, and *bla*_IMP_) detected by qPCR or GeneXpert® Carba-R from a LTCF was assessed over 16 months and compared to clinical infections. *bla*_KPC_, *bla*_OXA-48-like_, and *bla*_VIM_ were routinely detected in composite wastewater samples (6.4±0.8 log_10_ gene copy (GC)/ml (100% of 61 samples), 5.5±0.8 (98%), and 5.9±1.3 (34%), respectively), passive samples (8.2±0.5 log_10_ GC/g (31% of 55), 6.6±0.5 (96%), and 6.6±0.7 (31%)) and sewer biofilm (6.0±0.7 log_10_ GC/cm^2^ of pipe (100% of 17), 5.0±0.7 (100%), and 5.8±2.7 (24%)). Resistomes of wastewater and sewer biofilms differed, but both contained *bla*_KPC,_ *bla*_VIM,_ and *bla*_IMP_. Passive sampling may be a suitable alternative to composite sampling. Wastewater surveillance is a promising addition to carbapenemase monitoring.

## 1 Introduction

Antimicrobial resistant infections caused over 35,000 deaths in 2017 in the United States (1) and 1,270,000 deaths worldwide (2), with U.S. costs exceeding $13.5 billion (1). By 2050, antimicrobial resistance (AR) related deaths are projected to rise by 70%, to more than 208 million (3). Among the Centers for Disease Control and Prevention “Urgent Threats” are carbapenem-resistant Enterobacterales (CRE), responsible for ∼13,100 U.S. hospital-associated infections (1). Intermediate and long-term acute care facilities are particularly vulnerable to CRE due to high transmission risk and limited AR infection treatment options (4).

Routine patient surveillance for infection and colonization by AR pathogens is costly (5) and invasive (6). Monitoring hospital wastewater offers a non-invasive, cost-effective alternative that captures signals of AR from large patient populations, including colonized individuals. Multiple studies have attempted to correlate community wastewater antimicrobial resistance genes with infections or hospitalizations in the associated community (for recent reviews see Wang, Chu (7) and Wang, Han (8)). Wastewater surveillance at the hospital-level remains limited, but globally, hospital wastewater has been reported to have CRE and beta-lactamase (*bla*) genes such as *bla*_KPC_, *bla*_GES_, *bla*_IMP_, *bla*_NDM_, *bla*_OXA_, *bla*_TEM_, and *bla*_VIM_ with gene abundance ranging from 5 to 15 log_10_ gene copies (GC)/ml (9–14). Limited longitudinal studies have monitored hospital wastewater in Scandinavia (15, 16) and the United States (17) to establish antimicrobial resistance trends in wastewater and correlations to patient infections. For example, hospital wastewater collected monthly over 2-years in Sweden detected a *bla*_NDM_ producing CRE in wastewater that matched a patient isolate (15).

Carbapenem-resistant infections are a growing threat in healthcare settings, highlighting the need for practical surveillance methods to detect outbreaks early. Wastewater surveillance may be such a solution. However, there is concern that other factors may influence wastewater gene concentrations such as collection and processing methods. In addition, external factors such as bacteria absorption to, growth in, and sloughing from sewer biofilms may impact gene concentrations in wastewater. To address these concerns, we monitored five carbapenemase genes (*bla*_KPC_, *bla*_VIM_, *bla*_OXA-48-like_, *bla*_NDM_, and *bla*_IMP_) in wastewater from a post-acute care rehabilitation hospital (hereafter rehabilitation hospital) over 16 months. Samples were collected semi-weekly to semi-monthly using composite, passive, and biofilm-based methods and analyzed via quantitative polymerase chain reaction (qPCR), metagenomics, and culture. Sampling method, hold time, and storage temperature were evaluated to determine impact of these parameters on data reliability, with an emphasis on approaches suitable for inexpensive and simple collection, extraction, transport, and analysis for isolated facilities without the necessary nearby diagnostic laboratories. By examining temporal carbapenemase-producing (CP) gene abundance and establishing trendlines for composite, passive, and biofilm samples, we aimed to establish a baseline of carbapenemase genes for an in-patient facility. Once a baseline is established, new patient infections with carbapenemase producing organisms (CPOs) may be detected as CP gene increases in sewer samples.

## 2 Materials and Methods

### 2.1 Ethics statement

This study used de-identified patient data correlated with wastewater samples. The University of Utah Institutional Review Board determined the protocol to be exempt under Category 4 of 45 CFR 46.101(b) (protocol #00177673; exemption date: November 26, 2024). All samples were handled in accordance with biosafety precautions.

### 2.2 Site description and wastewater parameters

Samples were collected from a sewer serving a single rehabilitation hospital in Utah, USA, with no other wastewater sources contributing to the flow. With a staff of approximately 140, the 75-bed facility provides inpatient and outpatient care (600 out-patient clinical visits per month), and food services. Each source is estimated to contribute 48% (inpatient), 2% (outpatient), 14% (staff), and 37% (food services) of water use, based on estimates of water use by category (18). The daily visitor count is not known. Confirmation of wastewater travel time from various facility floors was done by flushing dye (Bright Dyes® tablet tracer dyes, Kingscote Chemicals, Miamisburg, OH) and visually monitoring wastewater in the sewer manhole for arrival of the appropriate dye. Water quality parameters (i.e., temperature, electrical conductivity, and pH) were collected at the same time as wastewater. Additional details are provided in the supplemental materials regarding the dye study and water quality parameter determination.

### 2.3 Limits of detection of AR genes and cultures in composite wastewater

CDC and FDA Antimicrobial Resistance Isolate Bank strains (see supplemental information; **Table S1**) were serially diluted in 40-ml of wastewater (n=5 for each dilution, with n=3 trials per gene) which contained no CP genes. Samples were then processed according to standard methods (see supplemental information, qPCR primers and probes in **Table S2**, positive controls in **Table S3**, qPCR reaction efficiencies in **Table S4**) and the viable culture and qPCR GC/ml were determined for each dilution. The lowest distinct dilution with a Ct above 35 was determined by ANOVA or ANOVA on ranks. Linear regression of Ct versus log CFU/ml was conducted for each trial separately and in aggregate for all trials along with the 95% confidence limits. Because wastewater used for this study already contained native CRO, we could not directly quantify the added antimicrobial-resistant strains on CHROMagar plates. Instead, we estimated the CFU/ml limit of detection from the initial culture concentrations added to each dilution.

### 2.4 Wastewater methods

#### 2.4.1 Sample collection methods

Three sampling methods (**Table 2**) were compared: composite wastewater (Hach AS950, Loveland, CO) collected over 24-hr, passive absorptive samples (tampons) typically deployed for 24-hr (n = 33) except when field conditions did not allow for a 24-hr collection (n = 29), and a sewer biofilm swab (EZ Reach™ Split sampler, World Bioproducts, Libertyville, IL) from the periodically wetted area of the sewer pipe above the flowing wastewater (see supplemental information).

#### 2.4.2 Sample processing methods

Bacteria were desorbed from passive samplers and biofilm swabs by vortex at max speed for 10 min in 1X phosphorus buffered saline (25 mL, 1X PBS, Thermo Fisher Scientific; Waltham, MA). All samples had a spike in recovery control (ZymoBIOMICS Spike-in Control I (Zymo), Zymo Research, Irvine, CA) added. Bacteria were precipitated by adding polyethylene glycol-6000 (PEG) and 0.9% of NaCl, shaking (80 RPM), centrifuging (typically 10 min, 31,400xg), and resuspension in 1.5 ml of 1X PBS. The resuspended pellet was subsampled for culturing [i.e., MacConkey I (Avantor; Radnor, PA) and CHROMagar KPC agar (DRG International; Springfield, NJ)], Cepheid GeneXpert® Carba-R analysis (Sunnyvale, California), and DNA extracted (19) for qPCR (20–22) and metagenomics (see supplemental information for additional details).

### 2.5 Comparison of sample volume, handling, and storage

The effect of concentrating 40 or 500-ml of sample for the gene abundance was investigated. The treatments compared 40-ml samples centrifuged at two different speeds and times compared to 500-ml samples (**Table 2**). These speeds and temperatures were tested with the understanding that not all laboratories have access to high-speed, temperature-controlled centrifuges. After centrifugation, nucleic acids were extracted from samples as described in supplemental materials. One-way ANOVA (SigmaStat, Systat Software Inc.; Palo Alto, California, v 14.5, with *P*<0.05) was used to assess significant differences among mean CP GC/ml when different water volumes were concentrated. If means were significantly different, multiple comparison tests were conducted using the Holm-Sidak or Dunn’s method.

The effect of sample handling conditions—including hold time, temperature, and processing stage—on gene abundance was evaluated in five treatment types, each with two or more replicates. Samples were concentrated using PEG centrifugation and resuspension in 1X PBS as described above. Samples processed immediately served as controls (**Table 2**). Controls were compared to samples held for different days and temperatures before concentration and nucleic acid extraction, and with and without preservation with 1:1 50% glycerol:10 mM 2-amino-2-(hydroxymethyl)propane-1,3-diol (Tris), pH 7.4. A general linearized model was used to determine the effect of sample handling and storage conditions on gene abundance as described in supplemental information.

### 2.6 Wastewater trends and alignment with facility patient testing data

The facility provided monthly average patient census, carbapenem days of therapy, numbers of patients screened for CRO, numbers of infections with CPO, percentage of patients that were independently performing all steps of toileting (i.e., contributing feces to the sewer) and showering, infected patient admit and discharge dates, and building water use.

Composite, passive, and sewer biofilm samples were collected weekly from January to December 2023 and then twice monthly from January to April 2024 (**Table 1**). All samples had CP gene composition analyzed by qPCR and Carba-R analysis. Total bacteria were determined by quantifying the conserved region of the 16S rRNA gene (aka Unibac primers)(20). The CP gene abundance by qPCR was always adjusted for Zymo recovery and log_10_ transformed. Further normalization of CP gene abundance by 16S rRNA GC or wastewater flow was conducted as described in the supplemental information.

**Table 1.**
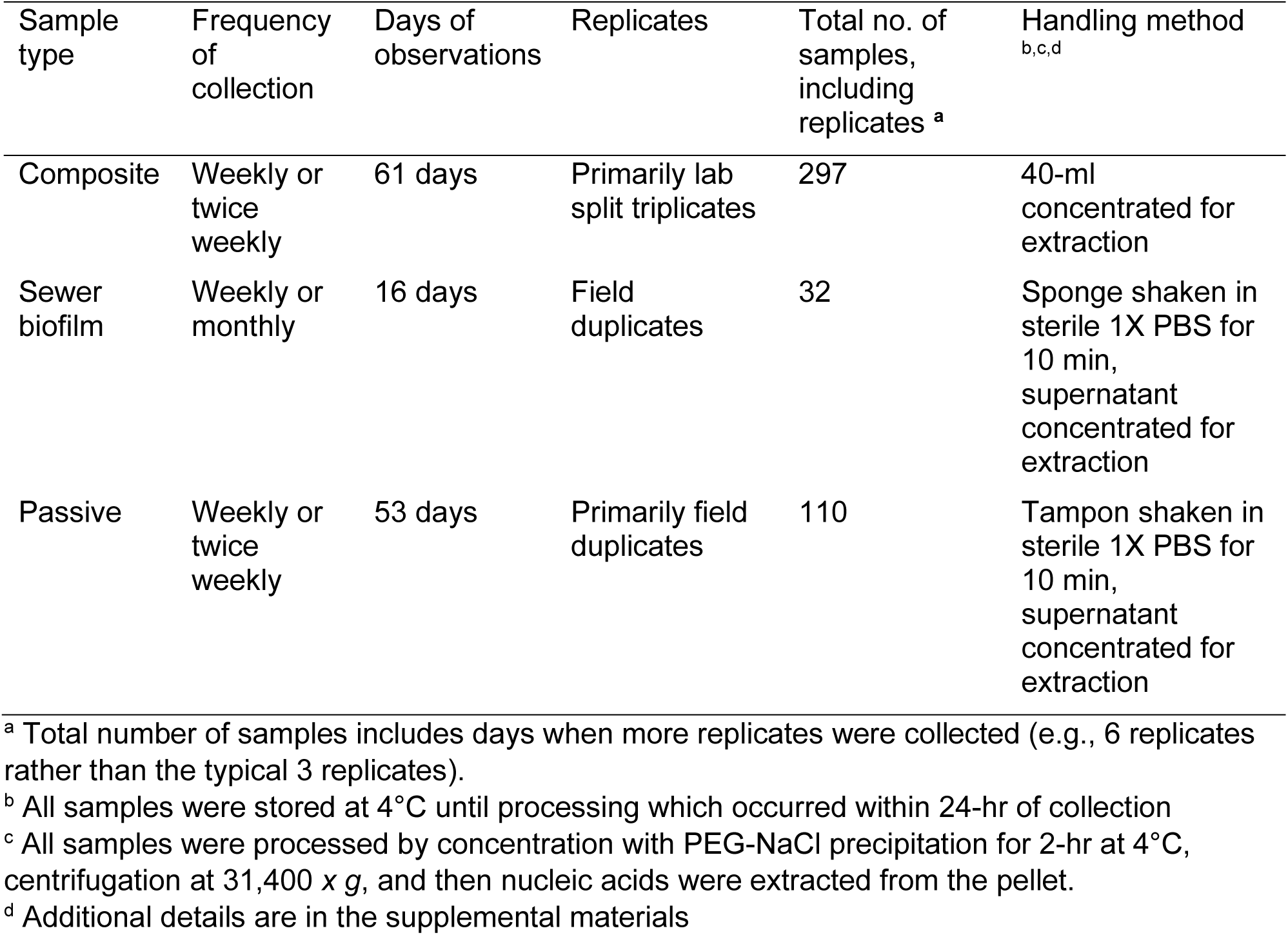
Sample types, frequency of collection, numbers of samples and handling methods for trend analysis over 16 months.

**Table 2.**
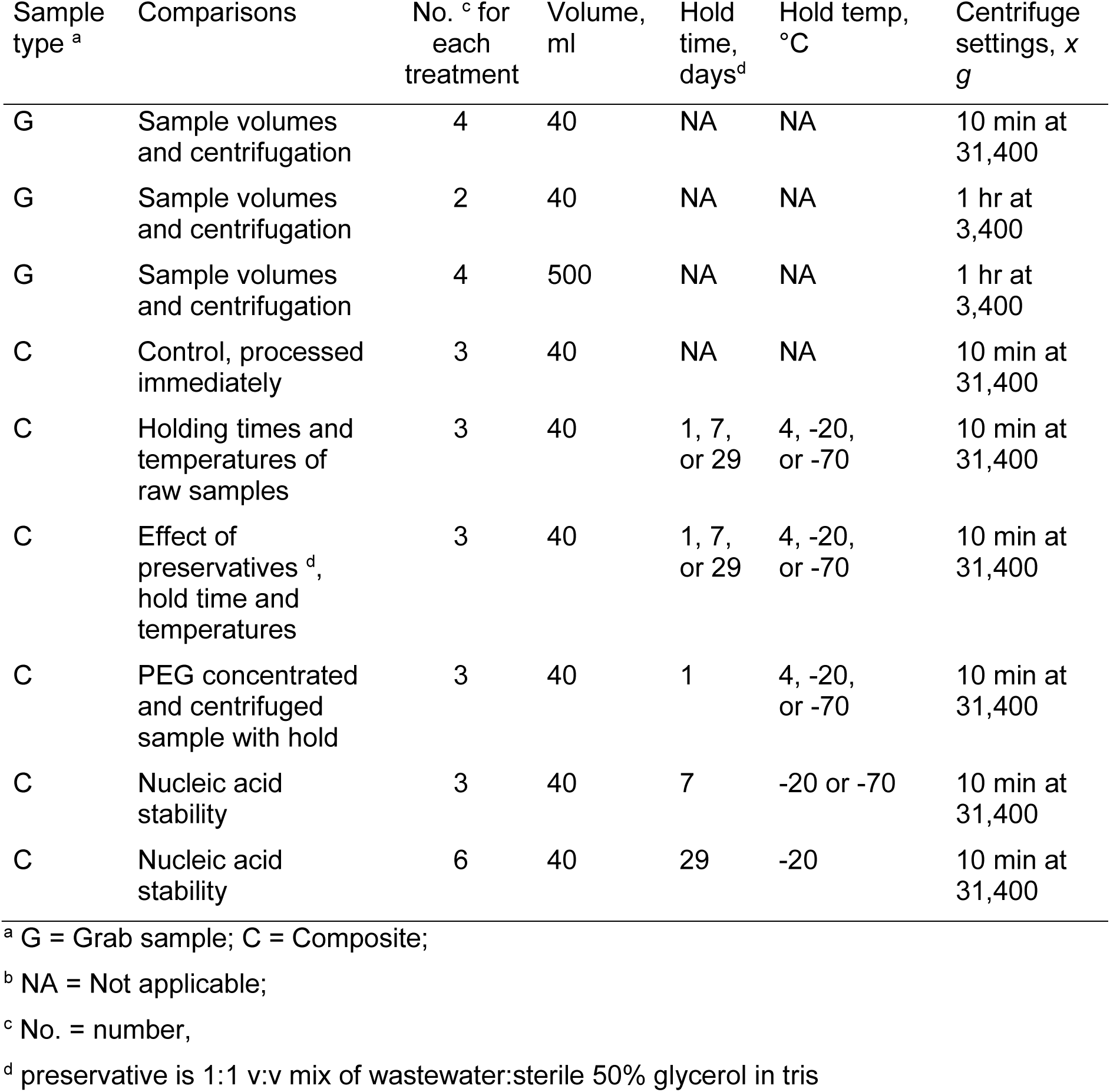
Sample handling methods for comparison of sample volumes, preservation method, hold time, hold temperature and centrifugation speeds.

Spearman rank order correlations of CP gene detections by method and multivariate linear regressions among ARG abundance were conducted in SAS (v9.4). A nonparametric regression technique was used to smooth the scatter plots of ARG over time using PROC LOESS in SAS (v9.4) with smoothing parameters from 0.1 to 0.25.

## 3 Results

### 3.1 Site description and wastewater parameters

The dye study and reviews of building drawings confirmed the manhole sampled received wastewater from all rehabilitation hospital floors. There was a 5- to 30-min travel time for the dye from sink drains closest to and farthest from the manhole suggesting 24-hour composite samples would capture recent patient stool and urine. Wastewater temperatures were 10.3 to 25.8°C (average x̅=20.8, n=55), conductivity was 1201 to 6330 µS/cm (x̅=1924, n=55), and pH was 6.4 to 9.0 (x̅=8.3, n=55).

### 3.2 Limits of detection of AR genes and cultures in composite wastewater

The detection limits of *bla*_NDM_, *bla*_VIM_, and *bla*_IMP_ in 40-ml of wastewater was 146 to 5486 GC/ml with average and standard deviations of three trials presented in **Table 3**. Variations in Ct observed for the different dilutions are shown in **Figure 1**. We found the lowest GC/ml determined by qPCR when extrapolated to the culture concentrations ranged from 25 to 337 CFU/ml with averages for *bla*_VIM_, *bla*_NDM_ and *bla*_IMP_ CFU/ml shown in **Table 3**. When the culture detection limit was estimated from linear regressions using all trials combined, the detection limits were lower (i.e., 17 to 216 CFU/ml), but had large 95% confidence limits (2.9 to 15437 CFU/ml, see **Table 3**). Detection limits of *bla*_KPC_ and *bla*_OXA-48-like_ were inconclusive due to the presence of those genes in wastewater used for dilution.

**Figure 1.**
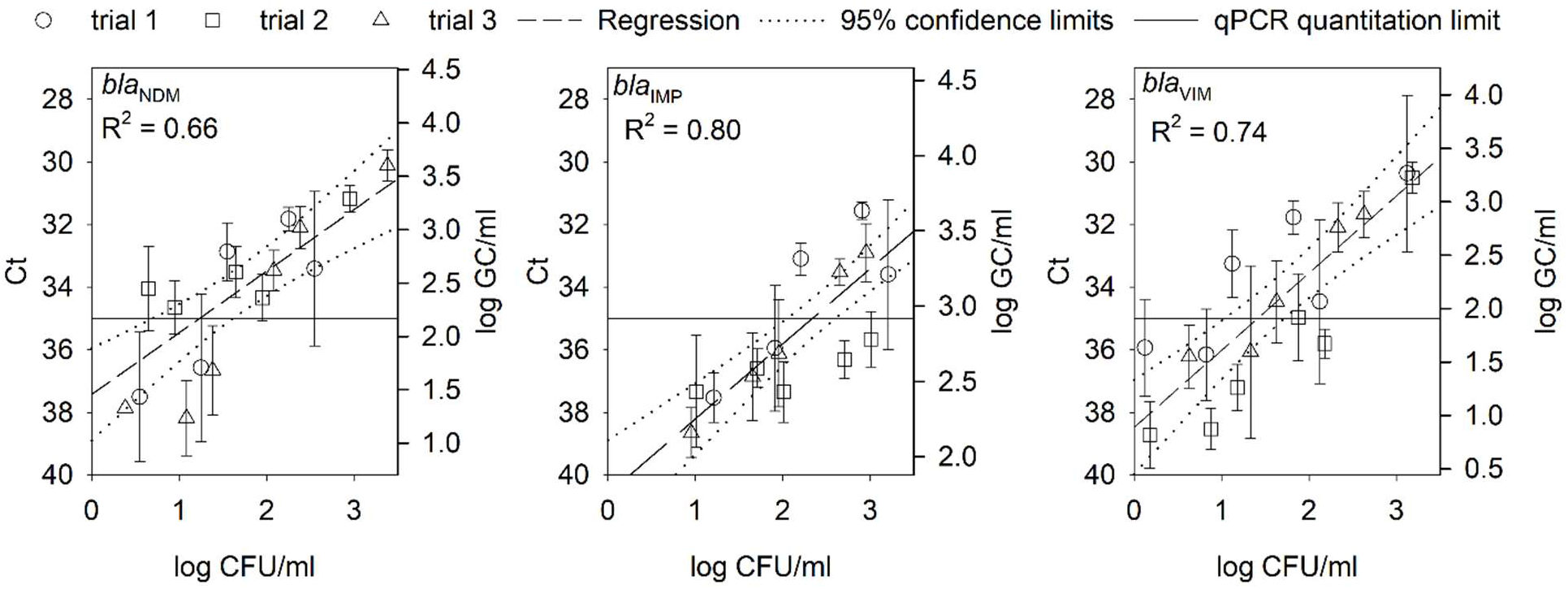
Mean and standard deviation of measured cycle thresholds (Cts), calculated log gene copies/ml and extrapolated log CFU/ml for the detection limit studies. Each point represents five replicates. Horizontal line represents a Ct cutoff of 35 cycles. All data points were included in the linear regression for each plot and the dashed lines represent the 95% confidence limits of the regression. The log CFU/ml was extrapolated from the initial isolate concentration that was then serially diluted by 6 log into each of 5 replicates. The log GC/ml were estimated from the Ct response from standard curves using either geneblock DNA or AR isolates as postive controls.

**Figure 2.**
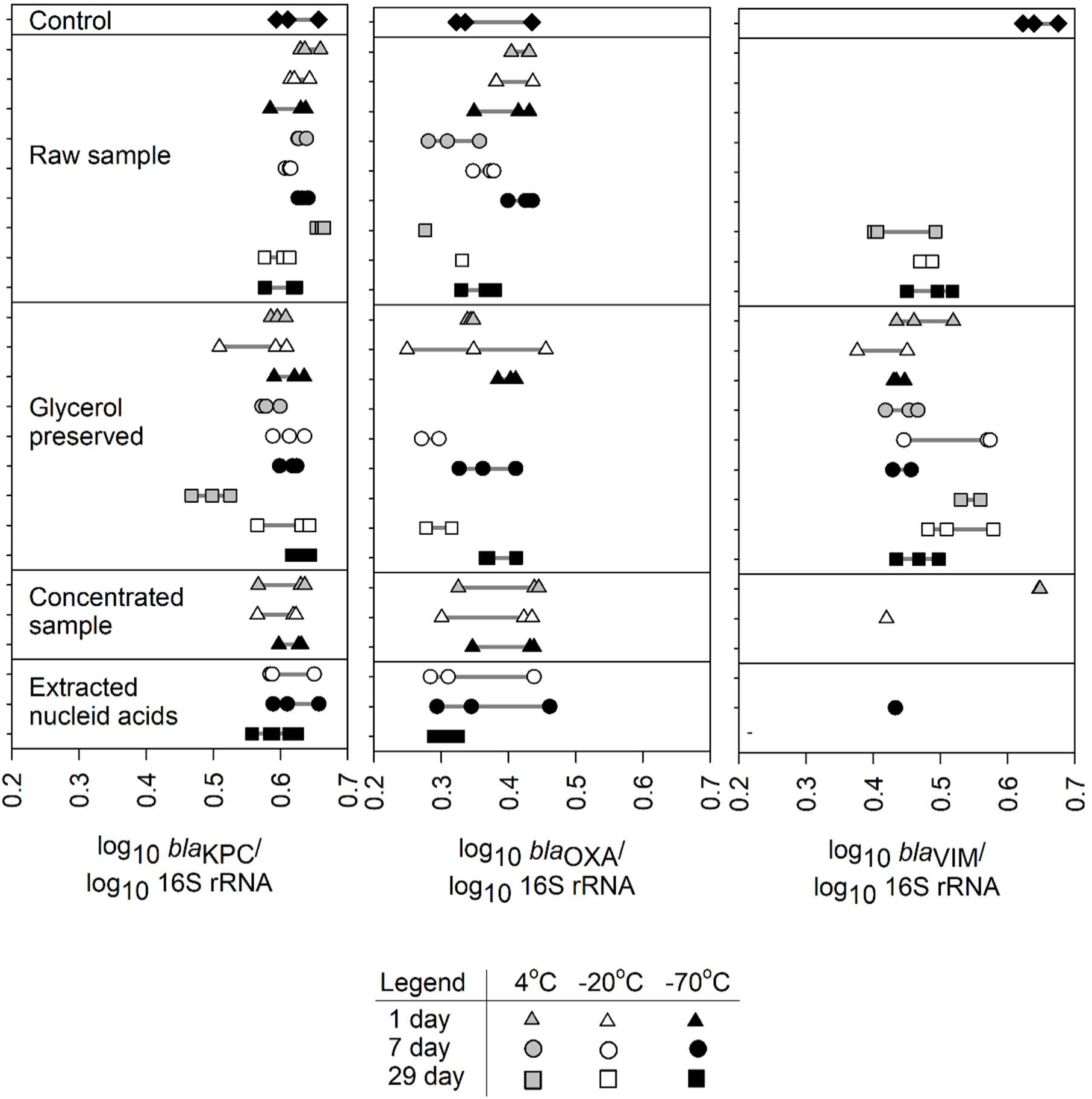
Holding time and preservation comparison for log_10_ *bla*_KPC_, *bla*_OXA-48-like_, and *bla*_VIM_ measured by qPCR in triplicate. All gene copies are normalized to log_10_ 16S rRNA gene. Horizontal grey lines were added to show the range of values for each treatment. Control sample DNA was extracted immediately upon receipt in laboratory. “Raw sample” are processed raw wastewater. “Glycerol preserved” are unprocessed samples preserved with 50% glycerol with Tris. Concentrated samples were PEG precipitated for 2-hr, centrifuged and the resulting pellet was resuspended and held at the listed temperature.

**Table 3.**
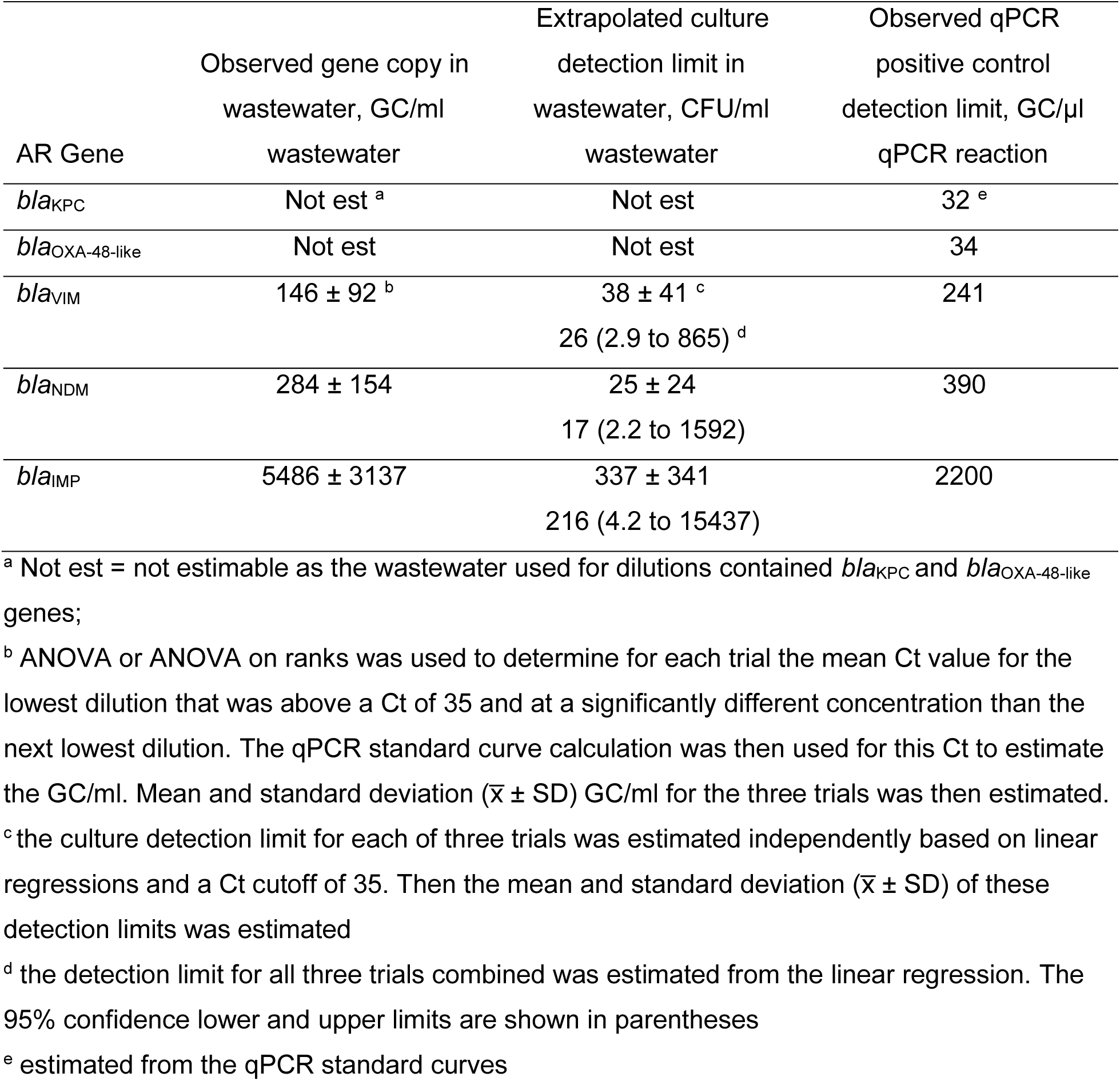
Limits of detection for isolates diluted in wastewater and detected by qPCR and culture based methods, and for qPCR positive controls diluted in PBS.

| AR Gene | Observed gene copy in wastewater, GC/ml wastewater | Extrapolated culture detection limit in wastewater, CFU/ml wastewater | Observed qPCR positive control detection limit, GC/μl qPCR reaction |
| --- | --- | --- | --- |
| *bla*<sub>KPC</sub> | Not est <sup>a</sup> | Not est | 32 <sup>e</sup> |
| *bla*<sub>OXA-48-like</sub> | Not est | Not est | 34 |
| *bla*<sub>VIM</sub> | 146 ± 92 <sup>b</sup> | 38 ± 41 <sup>c</sup><br>26 (2.9 to 865) <sup>d</sup> | 241 |
| *bla*<sub>NDM</sub> | 284 ± 154 | 25 ± 24<br>17 (2.2 to 1592) | 390 |
| *bla*<sub>IMP</sub> | 5486 ± 3137 | 337 ± 341<br>216 (4.2 to 15437) | 2200 |

<sup>a</sup> Not est = not estimable as the wastewater used for dilutions contained *bla*<sub>KPC</sub> and *bla*<sub>OXA-48-like</sub> genes;
<sup>b</sup> ANOVA or ANOVA on ranks was used to determine for each trial the mean Ct value for the lowest dilution that was above a Ct of 35 and at a significantly different concentration than the next lowest dilution. The qPCR standard curve calculation was then used for this Ct to estimate the GC/ml. Mean and standard deviation ( $\bar{x} \pm SD$ ) GC/ml for the three trials was then estimated.
<sup>c</sup> the culture detection limit for each of three trials was estimated independently based on linear regressions and a Ct cutoff of 35. Then the mean and standard deviation ( $\bar{x} \pm SD$ ) of these detection limits was estimated
<sup>d</sup> the detection limit for all three trials combined was estimated from the linear regression. The 95% confidence lower and upper limits are shown in parentheses
<sup>e</sup> estimated from the qPCR standard curves

### 3.3 Wastewater methods

#### 3.3.1 Comparison of composite, passive and biofilm sampling methods

The *bla*_KPC_, bla_VIM_ and *bla*_OXA-48-like_ GC normalized by water usage showed a strong correlation among composite wastewater and passive samples (**Table 4, Figure S2**, Spearman, *P*<0.05). In contrast, *bla*_NDM_ and *bla*_IMP_ were not correlated. In biofilm samples, *bla*_OXA_ was correlated with passive samples and with *bla*_KPC_. Inverse correlations between *bla*_KPC_ in biofilms and *bla*_VIM_ in passive samplers were observed, but only in 7 samples, thus decreasing the power of the analysis. Regardless of method, fewer gene concentrations were correlated when the data were not water usage normalized (**Table S5, Figure S3**). When normalized to facility water usage, sewer biofilm qPCR concentrations did correlate with composite wastewater samples.

**Table 4.**
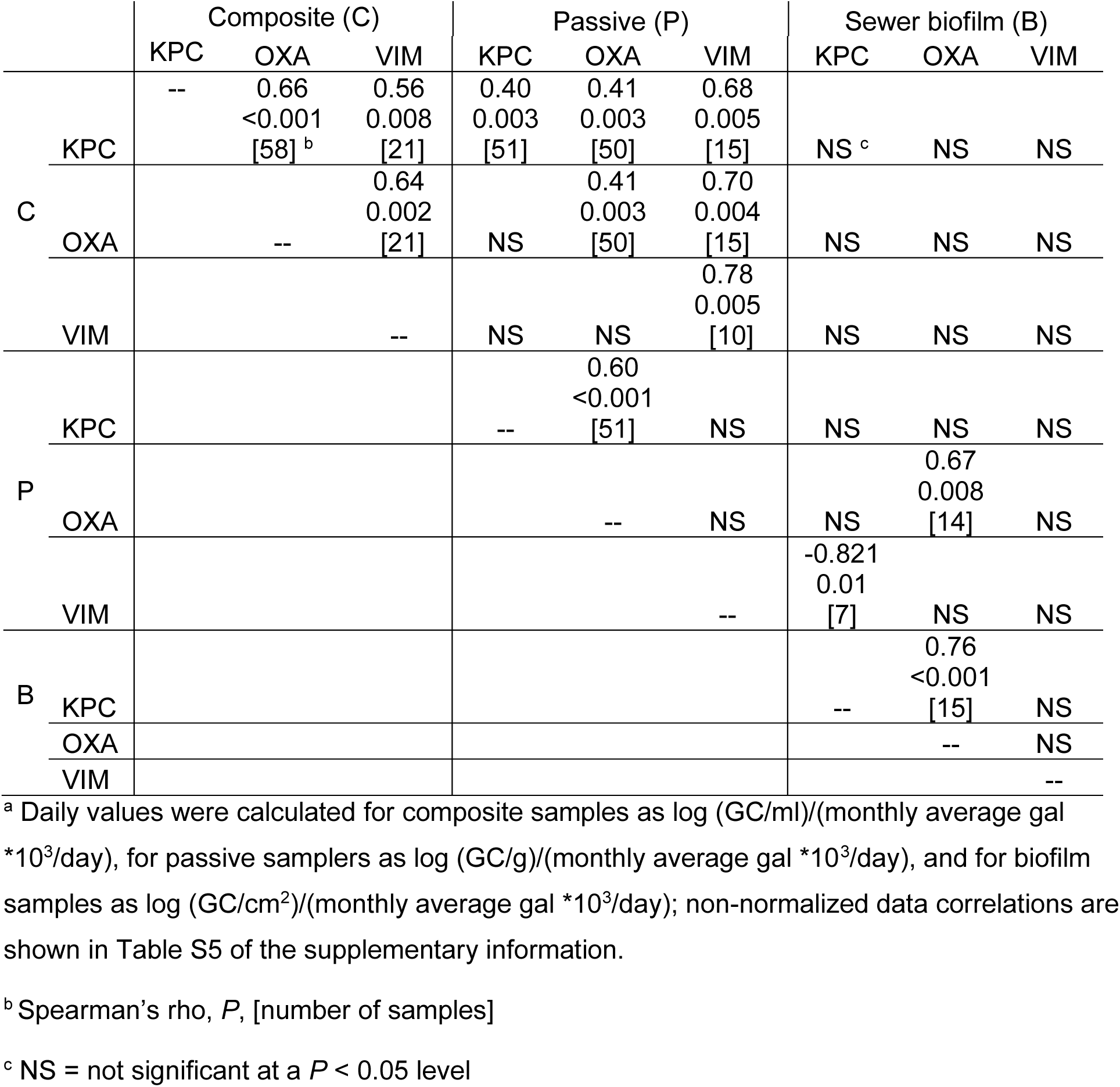
Spearman rank order correlations among building water usage-normalized^a^ *bla*_KPC_ (KPC), *bla*_OXA-48-like_ (OXA), and *bla*_VIM_ (VIM) log gene concentrations for composite, passive samplers, and sewer biofilm sample collection methods. *bla*_NDM_ and *bla*_IMP_ are not included as they were not correlated or only correlated in ≤ 5 fewer samples.

Deployment of passive samplers for longer than 24-hr (n=29) was shown to increase only total bacteria (i.e., conserved region of 16S rRNA gene) and *bla*_KPC_ (*P*<0.05) (i.e., 0.8- to 7-day deployment with 2.3 ± 1.5 days, average ± standard deviation), but R^2^ values were all less than 0.15 (**Figure S4**).

CHROMagar KPC and MacConkey I culture counts were correlated in passive or composite samples, for *E. coli*, *Pseudomonas* sp. and *Acinetobacter* sp. (**Tables S6** and **S7**). Contingency tables comparing sampling methods (**Table S8**) revealed that *bla*_KPC_ and *bla*_OXA-48-like_ genes were detected in most samples. Only 13 (3.9%) of 336 samples had *bla* genes detected in one sample type and not another by qPCR.

Metagenome sequencing revealed a wide range of antibiotic resistance genes (ARGs, **Figure S5**) including detection of *bla* genes in all sample types (**Figure S6**). Beta-lactamases and macrolide resistance genes had higher relative abundances in the biofilm sample (i.e., greater proportions of total sequences in that sample). Diversity indices for ARG (**Table S9**) show that grab and biofilm samples had the lowest richness, evenness, and Shannon index, suggesting that biofilms are dominated by a smaller number of highly abundant ARGs. A greater number of ARGs were detected in composite and passive samples (165-218 ARGs per metagenome) than the biofilm samples (155 ARGs).

All target *bla* genes were detected in the metagenomes of at least one sample (**Figure S5)**. Only *bla*_IMP-58_ variant was found in the metagenomes. *bla*_IMP_ was detected in the qPCR assay, but above the reportable threshold. Passive samples had more *bla*_KPC_ (**Figure S6**) compared to very low abundances (1-5 transcripts per million (TPM)) in the biofilm and grab metagenomes. *bla*_VIM_ was most abundant (14 TPM) in the biofilm metagenome. Neither *bla*_NDM_ nor any *bla*_OXA-48-like_ ARGs were detected in any metagenomes, despite detection by qPCR. We examined the relationship between metagenomic sequence coverage and qPCR abundance (**Figure S7**) finding a significant correlation for *bla*_KPC_ (R^2^=0.66). In contrast *bla*_VIM_ was only detected in one of three metagenomes where the qPCR abundance was 4 log_10_ GC. The *bla*_OXA-48-like_ and *bla*_NDM_ abundance was 2 to 6 log_10_ GC and <log_10_ 5 GC, respectively, but they were not detected in metagenomes. Therefore, the absence of *bla*_OXA-48-like_ and *bla*_NDM_ genes in the metagenomes is somewhat surprising but reasonable due to limits of detection for metagenomic sequencing.

Seven isolates from composite wastewater in January 2023 were sequenced and classified as *Citrobacter*, one was *E. coli* (serotype O8 + H19), and one *Providencia*. Six of 12 isolates from October 2023 were also *Citrobacter,* three *Comamonas*, and three *Aeromonas*. The *Aeromonas* and *Citrobacter* isolates had similar ARGs (**Figure S8**), including *bla*_KPC-2_, *bla*_KPC-3_, *bla*_OXA-2_, *bla*_TEM-1_, and *bla*_SHV_. The *E. coli* isolate genome includes *bla*_CTX-M-15_, *bla*_EC_, and *bla*_TEM-1_.

#### 3.3.2 Comparison of CP gene detection methods

Among all samples and collection methods there was close agreement between the Carba-R assay and qPCR detections (**Table S10-S12**), where 551 (85.4%) of 645 replicate CP gene assays were either both present or absent. *bla*_VIM_ was more frequently detected by Carba-R assay than qPCR with 76 (58.9%) of 129 samples Carba-R positive while qPCR was negative. *bla*_IMP_ was not detected by Carba-R, while 7% of samples (5 of 69) were qPCR positive.

#### 3.3.3 Comparison of composite wastewater sample volume, handling, and storage

Studies to evaluate the effect of wastewater (1) sample volume, (2) sample concentration method and (3) sample handling methods, on ARG abundance were conducted. In the first study of wastewater sampling volumes, we found no significant difference (Dunn’s method, *P*>0.05) in gene abundance among two 40-ml samples centrifuged at different speeds (31,400 *x g* or 3,400 *x g*) and times (10 or 60 min); therefore, these data were combined for further analysis. Overall, 40-ml samples had a significantly higher mean abundance compared to the 500-ml samples for *bla*_VIM_ (Holm-Sidak method, *P*<0.05), 16S rRNA gene, and *bla*_OXA-48 like_ (ANOVA on Ranks, *P*=0.02 and *P*=0.01, respectively), but not for *bla*_KPC_ (ANOVA, *P*=0.1) (**Figure S1**). Based on this, 40-ml samples, with centrifugation at 31,400 *x g* for 10-min was used herein.

Sample holding times (1, 7, and 29 days), holding methods (concentrated before storage or not) and preservation methods (raw sample or glycerol preserved) were compared to samples extracted and analyzed immediately to determine how sample handling influenced ARG abundance (**Figure 5**, **Table 5**). The type of sample preservation method (e.g., raw wastewater, 50% glycerol mix, concentrated, or DNA extracted (**Table 5**) generated the most variation in gene abundance. Specifically, the sample preservation method accounted for 24 to 42% of the variation in GC/ml. The highest recovery compared to the controls was observed in concentrated samples (57% to 121% average recovery of ARG among all temperatures and times) or extracted nucleic acid samples (59% to 94% recovery). Sample hold time prior to sample handling accounted for 25 to 38% of the variability. Hold time significantly influence ARG abundance (**Table 5**), thus shorter hold times are recommended.

**Table 5.**
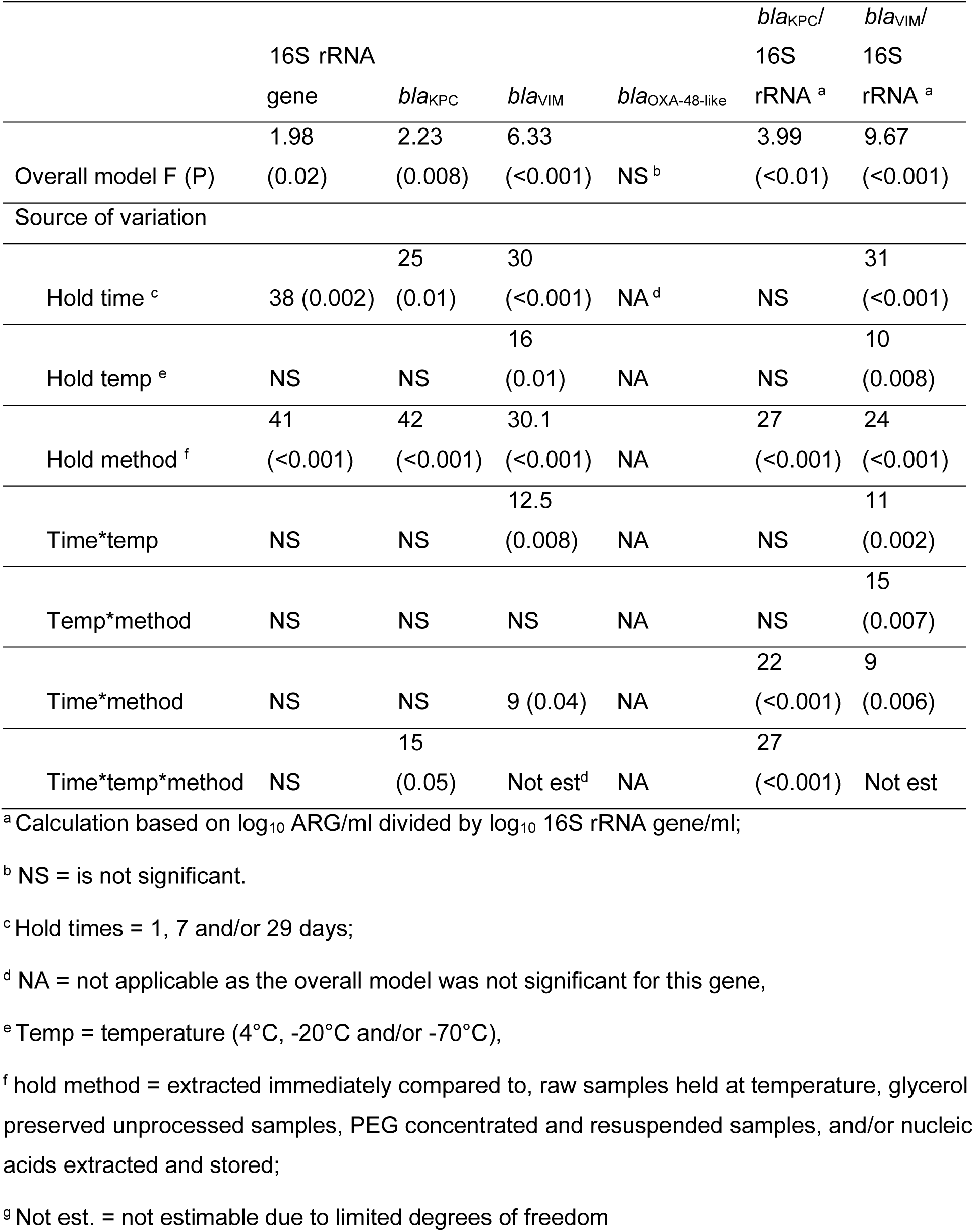
Effects of sample handling and storage conditions on gene abundance determined by qPCR. Source of variation is percent of total variation (*P*-value) of type 1 error for the general linearized model.

### 3.4 Wastewater CP gene trends and alignment with facility patient testing data

During the 16-month study, five CRO clinical infections were identified in four patients within the facility. These included: (infection 1 and 2) *Klebsiella (Enterobacter) aerogenes* and *Enterobacter cloacae* isolated from urine specimens collected nine days apart in April 2023 from the same patient; both were negative for CP genes; (infection 3) carbapenem-resistant *Pseudomonas aeruginosa* (CRPA) from a wound in April 2023, not tested for CP genes; (infection 4) CRPA from a urine specimen in December 2023, not tested for CP genes, and (infection 5) *Escherichia coli* and *Citrobacter freundii*, from a perianal screen in March 2024, not tested for CP genes. Notably, the patient with CRE identified in urine specimens in April 2023 used ostomy bags, which may not have been disposed of in the toilet. All other patients were recorded as toileting by nurses. Carbapenem antibiotic use in the facility during the observation period is in **Table S13**.

Over 16 months (**Table 6** and **7**) *bla*_KPC_ and *bla*_OXA-48-like_ were detected in >98% of composite and passive samples, but *bla*_VIM_, *bla*_NDM_ and *bla*_IMP_ were detected less frequently (35%, 10% and 3% in composite and 31%, 9% and 9%, in passive samples, respectively). All sewer biofilm samples contained *bla*_KPC_ and *bla*_OXA-48-like_ but only 24% of samples contained *bla*_NDM_ and *bla*_VIM_, and 18% contained *bla*_IMP_ (**Table 8**).

**Table 6.** Concentrations of CP genes in composite wastewater by month (mean ± standard deviation of log gene copies/mL) and frequency of CP gene detection (n, % positive) over 16 months.

| Mon-Yr<br>(n)<sup>a</sup> | *bla*<sub>KPC</sub> | *bla*<sub>OXA-48-like</sub> | *bla*<sub>VIM</sub> | *bla*<sub>NDM</sub> | *bla*<sub>IMP</sub> |
| --- | --- | --- | --- | --- | --- |
| 01-2023<br>(n=1) | 7.8<sup>b</sup><br>[1, 100%] | 6.3<br>[1, 100%] | 8.7<br>[1, 100%] | ND<sup>c</sup> | ND |
| 02-2023<br>(n=2) | 7.1±0.1<br>[2, 100%] | 6.2±0.4<br>[2, 100%] | 7.8±0.3<br>[2, 100%] | ND | ND |
| 03-2023<br>(n=4) | 7.3±0.3<br>[4, 100%] | 5.8±0.2<br>[4, 100%] | 7.3<br>[1, 25%] | ND | ND |
| 04-2023<br>(n=3) | 6.5±0.6<br>[3, 100%] | 4.9±0.5<br>[3, 100%] | ND | ND | ND |
| 05-2023<br>(n=8) | 6.9±0.5<br>[8, 100%] | 5.8±0.8<br>[8, 100%] | 6.1±0.6<br>[2, 25%] | ND | ND |
| 06-2023<br>(n=9) | 6.5±0.4<br>[9, 100%] | 5.4±0.5<br>[9, 100%] | 5.4±0.5<br>[2, 22%] | ND | ND |
| 07-2023<br>(n=7) | 6.3±0.6<br>[7, 100%] | 5.8±0.6<br>[6, 86%] | 4.2<br>[1, 14%] | ND | ND |
| 08-2023<br>(n=6) | 5.2±0.6<br>[6, 100%] | 5.2±0.7<br>[6, 100%] | 5.5±0.5<br>[2, 33%] | ND | ND |
| 09-2023<br>(n=4) | 6.1±0.8<br>[4, 100%] | 5.0±0.5<br>[4, 100%] | 5.2<br>[1, 25%] | ND | ND |
| 10-2023<br>(n=3) | 6.3±0.5<br>[3, 100%] | 5.4±0.1<br>[3, 100%] | 5.3±0.2<br>[2, 67%] | 4.6<br>[1, 33%] | ND |
| 11-2023<br>(n=4) | 6.1±0.4<br>[4, 100%] | 4.8±0.3<br>[4, 100%] | 4.8±0.2<br>[3, 75%] | 5.7<br>[1, 25%] | ND |
| 12-2023<br>(n=2) | 5.5±0.1<br>[2, 100%] | 4.4±0.2<br>[2, 100%] | 4.8<br>[1, 50%] | ND | ND |
| 01-2024<br>(n=2) | 5.6±0.2<br>[2, 100%] | 4.8±0.5<br>[2, 100%] | ND | ND | ND |
| 02-2024<br>(n=2) | 6.1±0.3<br>[2, 100%] | 5.7±0.3<br>[2, 100%] | 5.6±0.2<br>[2, 100%] | 5.1±0.6<br>[2, 100%] | ND |
| 03-2024<br>(n=2) | 5.8±0.9<br>[2, 100%] | 5.1±0.6<br>[2, 100%] | 4.8<br>[1, 50%] | 4.6±0.6<br>[2, 100%] | 6.5<br>[1, 50%] |
| 04-2024<br>(n=2) | 5.9±0.5<br>[2, 100%] | 5.0±0.4<br>[2, 100%] | ND | 5.3±0.4<br>[2, 100%] | 6.6<br>[1, 50%] |
| Totals |  |  |  |  |  |
| n = 61 | 61 | 60 | 21 | 8 | 2 |
| % positive | 100 | 98 | 34 | 13 | 3 |

<sup>a</sup> n = number of sampling dates in the month; <sup>b</sup> Concentrations without standard deviations indicate CP genes were detected in only one sample among multiple sampling dates in a month or only one sampling date in a month; <sup>c</sup> ND = not detected.

**Table 7.** Concentrations of CP genes collected with passive samplers by month (mean ± standard deviation of log gene copies/g) and frequency of CP gene detection (n, % positive) over 16 months.

| Mon-Yr<br>(n)<sup>a</sup> | *bla*<sub>KPC</sub> | *bla*<sub>OXA-48-like</sub> | *bla*<sub>VIM</sub> | *bla*<sub>NDM</sub> | *bla*<sub>IMP</sub> |
| --- | --- | --- | --- | --- | --- |
| 01-2023<br>(n=1) | 8.03<sup>b</sup><br>[1, 100%] | 6.8<br>[1, 100%] | 7.9<br>[1, 100%] | ND<sup>c</sup> | ND |
| 02-2023<br>(n=2) | 7.5±0.1<br>[2, 100%] | 5.9±0.1<br>[2, 100%] | 7.9±0.2<br>[2, 100%] | ND | ND |
| 03-2023 | Not tested | Not tested | Not tested | Not tested | Not tested |
| 04-2023<br>(n=1) | 9.0<br>[1, 100%] | 6.2<br>[1, 100%] | ND | ND | ND |
| 05-2023<br>(n=8) | 8.5±0.6<br>[8, 100%] | 7.0±0.7<br>[8, 100%] | ND | ND | ND |
| 06-2023<br>(n=9) | 8.3±0.5<br>[9, 100%] | 6.9±0.4<br>[9, 100%] | 6.8<br>[1, 11%] | ND | ND |
| 07-2023<br>(n=7) | 8.3±0.3<br>[6, 86%] | 6.5±0.2<br>[6, 86%] | ND | ND | ND |
| 08-2023<br>(n=7) | 7.9±0.3<br>[7, 100%] | 6.4±0.1<br>[6, 86%] | ND | ND | ND |
| 09-2023<br>(n=3) | 8.0±0.5<br>[3, 100%] | 6.4±0.4<br>[3, 100%] | ND | ND | ND |
| 10-2023<br>(n=3) | 7.9±0.2<br>[3, 100%] | 6.1±0.1<br>[3, 100%] | 6.4<br>[1, 33] | 5.4<br>[1, 33%] | ND |
| 11-2023<br>(n=4) | 8.1±0.4<br>[4, 100%] | 6.1±0.2<br>[4, 100%] | 6.0±0.2<br>[3, 75%] | 6.0<br>[1, 25%] | ND |
| 12-2023<br>(n=2) | 7.8±0.001<br>[2, 100%] | 6.3±0.2<br>[2, 100%] | 6.4±0.1<br>[2, 100%] | ND | ND |
| 01-2024<br>(n=2) | 7.9±0.03<br>[2, 100%] | 6.5±0.1<br>[2, 100%] | 6.3±0.1<br>[2, 100%] | ND | ND |
| 02-2024<br>(n=2) | 8.3±0.7<br>[2, 100%] | 6.8±0.5<br>[2, 100%] | 6.4±0.2<br>[2, 100%] | 5.1<br>[1, 50%] | 7.9±0.3<br>[2, 100%] |
| 03-2024<br>(n=2) | 8.2±0.7<br>[2, 100%] | 7.0±0.9<br>[2, 100%] | 6.2<br>[1, 50%] | 5.4<br>[1, 50%] | 7.8<br>[1, 50%] |
| 04-2024<br>(n=2) | 8.0±0.6<br>[2, 100%] | 6.7±0.2<br>[2, 100%] | 6.2±0.5<br>[2, 100%] | 6.1<br>[1, 50%] | 7.6±0.2<br>[2, 100%] |
| Totals |  |  |  |  |  |
| n = 55 | 54 | 53 | 17 | 5 | 5 |
| % positive | 98 | 96 | 31 | 9 | 9 |

<sup>a</sup> n = number of sampling dates in the month; <sup>b</sup> Concentrations without standard deviations indicate CP genes were detected in only one sample among multiple sampling dates in a month or only one sampling date in a month; <sup>c</sup> ND = not detected.

**Table 8.** Concentrations of CP genes observed in sewer biofilms by month (mean ± standard deviation of log gene copies/cm^2^) and frequency of CP gene detection (n, % positive) over 16 months.

| Mon-Yr<br>(n)<sup>a</sup> | *bla*<sub>KPC</sub> | *bla*<sub>OXA-48-like</sub> | *bla*<sub>VIM</sub> | *bla*<sub>NDM</sub> | *bla*<sub>IMP</sub> |
| --- | --- | --- | --- | --- | --- |
| 01-2023<br>(n= 1) | 4.6<sup>b</sup><br>[1, 100%] | 3.9<br>[1, 100%] | 6.9<br>[1, 100%] | ND<sup>c</sup> | ND |
| 02-2023<br>(n= 1) | 4.4<br>[1, 100%] | 3.9<br>[1, 100%] | 6.0<br>[1, 100%] | ND | ND |
| 03-2023<br>(n= 1) | 5.6<br>[1, 100%] | 3.5<br>[1, 100%] | ND | ND | ND |
| 04-2023 | Not tested | Not tested | Not tested | Not tested | Not tested |
| 05-2023<br>(n= 1) | 6.9<br>[1, 100%] | 5.7<br>[1, 100%] | ND | ND | ND |
| 06-2023<br>(n= 1) | 6.0<br>[1, 100%] | 5.0<br>[1, 100%] | ND | ND | ND |
| 07-2023 | Not tested | Not tested | Not tested | Not tested | Not tested |
| 08-2023<br>(n= 1) | 4.5<br>[1, 100%] | 4.0<br>[1, 100%] | ND | ND | ND |
| 09-2023<br>(n= 1) | 6.3<br>[1, 100%] | 5.6<br>[1, 100%] | ND | ND | ND |
| 10-2023<br>(n= 1) | 5.4<br>[1, 100%] | 4.5<br>[1, 100%] | ND | ND | ND |
| 11-2023<br>(n= 3) | 5.6±0.8<br>[3, 100%] | 4.2±1.1<br>[3, 100%] | ND | ND | ND |
| 12-2023<br>(n = 1) | 4.8<br>[1, 100%] | 4.1<br>[1, 100%] | 5.0<br>[1, 100%] | ND | ND |
| 01-2024<br>(n=1) | 5.0<br>[1, 100%] | 4.1<br>[1, 100%] | ND | ND | ND |
| 02-2024<br>(n = 1) | 5.3<br>[1, 100%] | 5.5<br>[1, 100%] | ND | 6.8<br>[1, 100%] | ND |
| 03-2024<br>(n = 1) | 5.1<br>[1, 100%] | 4.8<br>[1, 100%] | ND | 4.5<br>[1, 100%] | 6.5<br>[1, 100%] |
| 04-2024<br>(n = 2) | 4.3±0.7<br>[2, 100%] | 4.5±0.2<br>[2, 100%] | 4.2<br>[1, 50%] | 5.0±0.1<br>[2, 100%] | 6.3±0.2<br>[2, 100%] |
| Totals |  |  |  |  |  |
| n = 17 | 17 | 17 | 4 | 4 | 3 |
| % positive | 100 | 100 | 24 | 24 | 18 |

<sup>a</sup> n = number of sampling dates in the month; <sup>b</sup> Concentrations without standard deviations indicate CP genes were detected in only one sample among multiple sampling dates in a month or only one sampling date in a month; <sup>c</sup> ND = not detected.

Abundance of *bla*_KPC_ and *bla*_OXA-48-like_, normalized for facility water usage, increased after April 2023 when CRE and CRPA infected patients were present (**Figure 3A, 3B, 3D,** and **3E**). A general increase in *bla*_KPC_ and *bla*_OXA-48-like_ abundance was also observed in the last four months of the study period (**Figure 3A, 3B, 3D,** and **3E**), although not attributable to a patient infection. The *bla*_VIM_ abundance generally decreased over the study period for flow-normalized and non-normalized data (**Figure 3G, 3H, 3I, and S9**).

**Figure 3.**
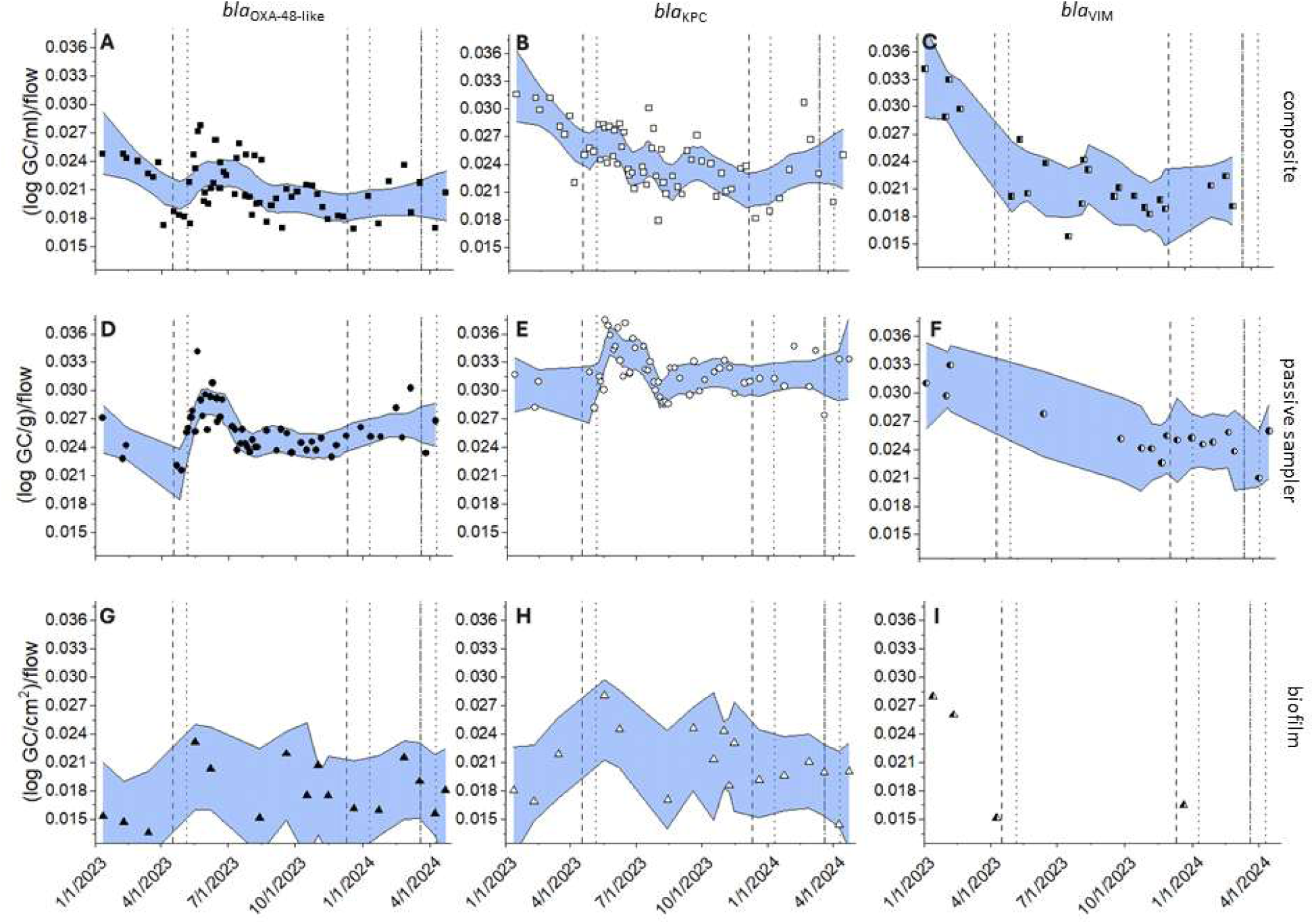
Gene copies (GCs) of *bla*_OXA48-like_ (left column, closed symbols), *bla*_KPC_ (center column, open symbols), and *bla*_VIM_ (right column, half-filled symbols) in wastewater composite collected by autosamplers (top row, circles), passive samplers (middle row, squares), or swabs of sewer biofilm (bottom row, triangles), from January 2023 to April 2024. Values are normalized by recovery control and water usage. Non parametric regression smoothing factors range from 0.1 to 0.25. Dash (admit date) and dotted (discharge date) vertical lines indicate periods when patients infected with CRE or CRPA were in the facility including *Klebsiella aerogenes*, *Enterobacter cloacae* and *Pseudomonas aeruginosa* in period 1 (April, 2023), *P. aerugniosa* in period 2 (December 2023) and *Escherichia coli* and *Citrobacter freundii* in period 3 (March 2024). The CRE infections in April 2023 and March 2024 were not CPO based on PCR. None of the CRPA were tested for CP genes.

## 4 Discussion

This study demonstrated the potential for wastewater-based surveillance to monitor CP genes from hospitals. Over 16 months, three carbapenemase genes, *bla*_KPC_, *bla*_VIM_, and *bla*_OXA-48-like_, were consistently detected by qPCR and Cepheid Carba-R assays in each of the three collection methods. Among sampling strategies, the composite and passive sampling consistently outperformed biofilm sampling in terms of gene richness and AR gene detection frequency. Passive samplers also yielded higher concentrations of carbapenemase genes, possibly due to prolonged absorption of bacteria and extracellular DNA. The sewer biofilm gene concentrations when normalized to building water usage was correlated with wastewater concentrations suggesting the sewer biofilm may influence the wastewater CP gene abundance, but additional studies are needed to determine the relationship. While all sample types contained *bla*_KPC_ and *bla*_OXA-48-like_, less frequently detected genes such as *bla*_VIM_, *bla*_NDM_, and *bla*_IMP_ were more variable across sampling methods. The Carba-R assay is known to generate a negative IMP result even when IMP-7, -13 or -14 gene sequences are present (24). The Carba-R assay was more likely to detect *bla*_VIM_ than qPCR but the results were not quantitative. Differences in gene abundance between sewer biofilms and composite samples may reflect shedding from colonized patients, environmental persistence, or microbial community dynamics.

Sample handling conditions significantly affected gene abundance measurements. Gene recovery was highest when 40 ml of sample was processed using a higher centrifuge speed compared to 500 ml with lower centrifugation speeds. Recovery decreased with prolonged storage or higher temperatures unless samples were preserved with glycerol. These findings emphasize the importance of standardized, practical protocols (25). The results show that passive sampling is as effective as composite sampling, and storing raw samples at 4°C up to 29 days was comparable to immediate processing. This finding is especially important for resource limited settings as these facilities can collect and store a passive sample prior to processing in lieu of composite samples processed the next day.

Temporal trends in wastewater gene abundance were unable to be correlated with CP genes in patient colonizing isolates due to limited infections during the sampling period. Due to limited infections in the facility during the study period, longer term sampling is required to evaluate the utility of wastewater surveillance as a complement to hospital AMR surveillance methods. Two critical points require further consideration: (1) wastewater results should be interpreted considering AR bacteria reservoirs like sinks and sewer biofilms, which have been found to harbor CP genes herein (**Figure 4**) and CRE as reported elsewhere (26, 27) and (2) CRE or CP gene sources in addition to infected patients such as colonized patients, hospital staff, outpatients, and visitors. Additional research is needed to investigate the contribution of sewer biofilms to wastewater from hospitals as these biofilm signals may confound temporal gene signals. Identifying the non-patient contribution of CRE or CP genes to wastewater will also be critical for successful surveillance efforts.

In conclusion, wastewater sampling—particularly composite and passive methods—combined with sensitive molecular detection may be a practical, non-invasive approach for monitoring carbapenemase genes in healthcare facilities. Passive samplers may be suitable in low resource settings or situations where space constraints or sewer access are concerns. When wastewater sampling is integrated with clinical data, it remains an open question whether such surveillance may enhance outbreak detection, support infection control, and aid antibiotic stewardship, and additional study is needed.

## Acknowledgements

The authors acknowledge the valuable assistance of Sophia Reyes, Nathan Hatten, Aspen Dalby, Daniel Jeun, Katherine Reilly, Vivian Marcoux, and the Utah Public Health Laboratory.

## Financial support

This work was supported by CDC contract #200-2021-12774, Safety and Healthcare Epidemiology Prevention Research Development (SHEPheRD) 2022 Domain 1-A004: Wastewater surveillance approaches for antimicrobial resistant genes and organisms in healthcare settings within the Western U.S. Region, Jennifer Weidhaas, Ph.D. (Civil and Environmental Engineering), Principal Investigator. This work was also supported in part by a first year fellowship to E. Warren from the University of Utah Global Change and Sustainability Center.

## Disclaimer

The conclusions, findings, and opinions expressed by authors contributing to this journal do not necessarily reflect the official position of the US Department of Health and Human Services, the Public Health Service, the Centers for Disease Control and Prevention, or the authors’ affiliated institutions. The use of trade names is for identification only and does not imply endorsement by the Public Health Service or by the United States Department of Health and Human Services.

## Author Contributions

<u>Ean Warren:</u> Data Curation, Formal Analysis, Investigation, Methodology, Visualization, Writing-original draft preparation. <u>Jim VanDerslice:</u> Conceptualization, Funding Acquisition, Project Administration, Writing-review and editing. <u>Scott Benson:</u> Conceptualization, Funding Acquisition, Data Curation, Writing-review and editing. <u>William Brazelton:</u> Conceptualization, Data Curation, Formal Analysis, Writing-review and editing. <u>Windy Tanner:</u> Conceptualization, Funding Acquisition, Methodology, Writing-review and editing. <u>Amanda Lyons:</u> Conceptualization, Funding Acquisition, Writing-review and editing. <u>Florence Whitehill:</u> Conceptualization, Funding Acquisition, Writing-review and editing. <u>Angela Coulliette-Salmond:</u> Conceptualization, Funding Acquisition, Writing-review and editing. <u>Sydney Fusco:</u> Data Curation, Investigation. <u>Jennifer Weidhaas:</u> Conceptualization, Formal Analysis, Funding Acquisition, Methodology, Project Administration, Supervision, Visualization, Writing-review and editing.

## Notes

### Competing Interest Statement

The authors have declared no competing interest.

