## Supplemental materials for "Evaluation of Detection Methods for Wastewater Surveillance of Antimicrobial-Resistant Bacteria from Healthcare Facilities"

^c^ University of Utah, Department of Family and Preventive Medicine,
^i^ United States Public Health Service, Rockville, MD, 20853

^k^ University of Utah, Department of Civil and Environmental Engineering,

* Corresponding author

[Table S6. Bacterial concentrations [log10(colony forming units/100 ml)] determined by CHROMagar KPC and MacConkey I. ^a^	19](#_Toc209361704)

[Figure S9. Gene copies (GCs) of *bla*_OXA‑48-like_ (left column, closed symbols), *bla*_KPC_ (center column, open symbols), and *bla*_VIM_ (right column, half-filled symbols) in wastewater collected by autosamplers (top row, circles), passive samplers (middle row, squares), or swabs of sewer biofilm (bottom row, triangles), from January 2023 to April 2024. Non parametric regression smoothing factors range from 0.1 to 0.25. Dash and dotted vertical lines indicate periods when patients infected with CRE or CRPA were in the facility including *Klebsiella aerogenes*, *Enterobacter cloacae* and *Pseudomonas aeruginosa* in period 1 (April, 2023), *P. aerugniosa* in period 2 (December 2023) and *Escherichia coli* and *Citrobacter freundii* in period 3 (March 2024). The CRE infections in April 2023 were not CPO based on PCR. The CRE infection in March 2024 was not tested for CP genes by PCR. None of the CRPA were tested for CP genes.	31](#_Toc209361721)

### Materials and Methods

#### Wastewater flow and water-quality analysis

Water usage was determined from water meter readings contained in monthly water bills that were provided by the facility to the research team. Wastewater temperatures were determined at the time of sample collection using a Raynger MX4PU noncontact, infrared thermometer (Raytek; Santa Cruz, California). Conductivity and pH were collected in the field or in the laboratory with a sub-sample of the composite sample. In either case, the conductivity and pH were measured using a calibrated labForce TS-PC200 (Thomas Scientific; Swedesboro, New Jersey) portable pH and conductivity meter with separate pH (PH-02) and conductivity (CC-01) probes. Analyses were performed within 2 hours of wastewater collection.

#### Sample collection methods

A Hach AS950 autosampler (Hach; Loveland, Colorado) was used to composite wastewater over 24 hours, consisting of 100 to 200 ml collected every 15 to 20 minutes into a single carboy. One liter of the composite was retained for analysis. All composite samples were split at the lab into triplicate 40 ml subsamples. Before each sample collection event, the autosampler sampling tubing was sterilized by pumping 1.5 L of fresh 10% bleach solution through the tubing, followed by flushing with 1.5 L wastewater.

Passive samples, also called Moore samples (1), consisted of sterile tampons (Equate brand tampons, size super; Tampax brand tampons, size super; and OB brand tampons, size regular). The average absorbency and mass were determined from 10 tampons from each brand. Overall, tampons in this study retained 12 to 17 ml of wastewater. Equate tampons average mass were 1.0 g; Tampax tampons were 1.5 g; and OB were 1.6 g. Duplicate tampons were placed in the sewer for at least 24 hours to collect wastewater. Retrieved tampons were placed into sterile centrifuge tubes containing 25-ml sterile 1X phosphate-buffered solution (PBS, Thermo Fisher Scientific; Waltham, Massachusetts). Between 12 January 2023 and 15 June 2023, Equate tampons were used while between 21 June 2023 and 27 July 2023, super-size Tampax brand tampons were used. All other sampling events after 27 July 2023 used OB tampons.

Biofilm samples were collected by wiping both sides of a sponge (EZ Reach™ Split) sampler (World Bioproducts; Libertyville, Illinois) over 40 cm^2^ of the sewer pipe above the water line in the periodically wetted area. Most sewer pipes from buildings are 6 to 8 inch (15.24 to 20.32 cm) in diameter. The wastewater in sewer pipes is typically flowing at a depth between 0.25 and 1 inch in depth or less than 1/3 of the depth of the pipe. With periodic flushing from a building, this water level will rise and then fall (aka periodically wetted). We sampled the area of the sewer pipes above the currently flowing wastewater line in the area that is periodically wetted. The sponge field duplicates were collected by aseptically sectioning and placement into 50-ml centrifuge tubes containing 25-ml sterile 1X PBS. Grab samples were collected in a sterile, 1-L polypropylene sample bottle.

#### Sample handling methods

Replicate wastewater samples, both composite and grab, were split into 40 ml triplicates. ZymoBIOMICs Spike-in Control I (Zymo Research; Irvine, California) containing *Imtechella halotolerans* and *Allobacillus halotolerans*, was added to obtain 1.25 x 10^6^ GC/μl. Then 10% polyethylene glycol-6000 (PEG) and 0.9% of NaCl were added. Passive or biofilm swabs in 25 ml 1X PBS were mixed using a vortex for 10 minutes to release any bacteria into the supernatant before removal of the tampon or sponge pieces. Then 10% PEG, 0.9% NaCl, and Zymo spike were added to the 25 ml retained solution. All sample types containing PEG/NaCl/spike were then shaken at 4°C for 2 hours followed by centrifugation at 31,400 *x g* for 10 min at 4°C. The supernatant was removed and the pellet was resuspended in 1.5 ml of sterile 1X PBS. The resuspended pellet was subsampled for culturing, Cepheid GeneXpert® Carba-R analysis (Sunnyvale, California), and DNA extracted for qPCR and metagenomics.

The 16S rRNA concentration of the spike-in is 1 x 10^7^ gene copies/μl (Zymo Research Spike-in Control I (Zymo)). Spike-in volume addition ranged from the smallest amount of 7 μl in 6.5 L (i.e., 1 x 10^4^ gene copies/μl) to 5 μl in 40 ml (i.e., 1.25 x 10^6^ gene copies/μl final concentration).

For centrifuge tubes that contained passive or biofilm swab samples in 25 ml 1X PBS, the tubes were first mixed by placing them horizontally on a vortex for 10 minutes to release any bacteria into the supernatant. Then, the tampons or sponge pieces were removed. Ten percent polyethylene glycol-6000 (PEG), 0.9% sodium chloride and the Zymo spike were added to the retained solution in the centrifuge tubes. Unless otherwise noted, all sample tubes were shaken at 4°C for at least two hours before centrifugation at 31,400 *x g* for 10 minutes at 4°C. The supernatant was poured off, retaining the pellet, then 1.5 ml 1X sterile PBS was added and the pellet was resuspended. The 1.5 ml of resuspended, concentrated pellet was subsampled for culturing, Cepheid GeneXpert Carba-R analysis, and DNA extraction for qPCR and metagenomics.

#### Culturing methods

Concentrated suspended samples (subsample of the 1.5 ml described above) were analyzed by plating on MacConkey I (Avantor, Randor, Pennsylvania) and CHROMagar KPC agar (DRG International, Springfield, New Jersey). The suspended pellet was serially diluted ten-fold into 1X PBS. Dilutions were plated on the MacConkey I or CHROMagar KPC plates and incubated at 37°C overnight for 18 to 24 hours. Colonies were counted, picked, and streaked into quadrants on tryptic soy broth plates with 5% defibrinated sheep’s blood (Thomas Scientific, Swedesboro, New Jersey) and agar (molecular genetics grade from Fisher Scientific, BP1423-500). An antimicrobial susceptibility disk containing 10 μg meropenem was placed in each streaked region and the plates were incubated at 37°C overnight. After incubation, the diameter of the affected bacterial lawn was measured. Picked colonies were incubated in centrifuge tubes with tryptic soy broth at 37°C for 16 to 72 hours. An equal amount of sterile 50% glycerol in 10 mM 2-Amino-2-(hydroxymethyl)propane-1,3-diol (Tris) was added as a preservative and the cultures were stored at ‑80°C. The colony forming units (CFU) per 100 ml of wastewater or PBS from the MacConkey I or CHROMagar KPC plates were recorded by colony color. Numbers of white and opaque colonies grown on CHROMagar KPC plates were combined due to the similarity of the two colony types.

#### Cepheid GeneXpert Carba-R method

The Cepheid GeneXpert Carba-R test detects the present or absence of *bla*_KPC_, *bla*_VIM_, *bla*_OXA‑48_, *bla*_NDM_, and *bla*_IMP_ carbapenemase genes. Samples were kept on ice packs for transportation to the laboratory. Fifty μl of resuspended pellets were added to the Cepheid Sample Reagent bottle and analyzed using the Cepheid standard protocol. All samples prior to 10 May 2023 were run individually (n=96 individual samples). The remaining replicates were combined in equal volumes before analysis to save resources and costs (n=241 sample lab splits (i.e., triplicate composite, duplicate passive, and duplicate biofilm) combined into 99 samples tested after May 2023). As per manufacturer’s method, the sample reagent bottle was mixed using a vortex and approximately 1.7 ml of the sample was added to the Carba-R cartridge using the transfer pipette provided by Cepheid. The cartridge was put into the GeneXpert instrument for analysis. The Xpert-Carba-R results are reported as only “detected” or “not detected” for the five carbapenemase genes tested. However, the instrument also reports the Ct values for each gene target which was recorded and tracked during data collection.

**Nucleic acid extraction**

DNA was extracted from 0.5 ml concentrated resuspended pellets using Lysing Matrix E bead beater tubes (MP Biomedicals; Irvine, California). The rest of the method generally followed a previously published manual extraction protocol (2). Briefly, samples were lysed with an equal mixture of cetyltrimethylammonium bromide (CTAB) and phenol-chloroform-isoamyl alcohol, pH 8 for 10 minutes while mixing using a vortex at setting 8. The samples were cleaned of phenol with chloroform:isoamyl alcohol (24:1). The PEG-precipitation was done for two hours at room temperature in the dark followed by centrifugation at 17,000 *x* *g* for 15 minutes. The pellet was washed with 100 μl 70% ice-cold ethanol then centrifuged at 17,000 *x* *g* for another 15 minutes. The nucleic acid pellet was dissolved in 50 μl 10 mM Tris buffer. Extracted nucleic acids were quantified using UV detection at 260 and 280 nm to determine approximate DNA concentrations and purity.

#### Quantitative Polymerase Chain Reaction for Carbapenemase Genes

A QuantStudio™ 3 (Applied Biosystems) was used for qPCR assays (3-5). Primer and probe sequences, and concentrations are found in **Table S2** while positive control sequences are found in **Table S1.** KiCqStart Probe qPCR ReadyMix (Sigma Aldrich; St. Louis, Missouri) was used for *bla*_VIM_ and *bla*_OXA‑48-like_ assays, while TaqMan™ Fast Advanced Master Mix (Thermo Fisher Scientific) was used for all other assays. Standard curves were created using 10-fold dilutions of the positive control. Efficiencies ranged from 88.9 to 119.0% (**Table S4**). Gene copy calculations and thermocycler conditions are shown in the supplemental material. The Ct cutoff for qPCR sample analyses was 35 cycles.

The gene copies (GCs) obtained by qPCR were normalized by volume for composite samples, by mass for passive samples, or by area swabbed for biofilm samples.

$$\frac{\mathrm{GC}}{ml wastewater}=average GC concentration*dilution*DNA diluent volume/wastewater volume$$

$$\frac{\mathrm{GC}}{g collector}=average GC concentration*dilution*DNA diluent volume/collectordry mass$$

$$\frac{\mathrm{GC}}{\mathrm{cm}^{2}\mathrm{swabbed}}=average GC concentration*dilution*DNA diluent volume/area swabbed$$

Typically, the passive sampler (i.e., tampons) weighed 1.0 to 1.6 g (dry weight). While passive samplers were routinely deployed in duplicate, occasionally one of the tampons was lost. In that case the remaining tampon sampler was cut in half in the field and the two pieces were used as field duplicates. The mass of the collector was then assumed to be half the usual mass when calculating GC/g collector.

The qPCR units for the different sample types included log gene copies/ml wastewater for the autosampler and grab samples, log gene copies/g for passive samplers, and log gene copies/cm^2^ of sewer pipe for biofilms. Comparison among different sample types (i.e., wastewater, passive sampler, or biofilm) with different units was conducted using either gene copies (GC) divided by 16S rRNA concentrations or GC adjusted for extraction recovery using Zymo spike control.

$$Measured Zymo concentration=Measured Zymo concentration by qPCR*dilution$$

In either case, the GC/unit concentration was divided by 16S rRNA concentration or Zymo fraction recovered using qPCR results for the same sample replicate. All log transforms are base 10.

#### Metagenomic Analysis and Sequencing

Metagenome sequencing of the different types of sample collection methods was conducted in the first two months of the project period to evaluate differences among the microbial communities collected by each method. Metagenome sequences of 7 wastewater samples were obtained, including 3 composite samples (Autosample1, 2, and 3), a Biofilm sample, a Grab sample, and 2 passive samples (Moore1 and 2). Autosample1, Biofilm, Grab, and Moore1 samples were collected on January 12, 2023. Autosample2 was collected on January 24, 2023. Autosample3 and Moore2 samples were collected on October 17, 2023. Metagenomic sequencing was conducted at Utah Public Health Laboratory with an Illumina NextSeq platform (P2 flow cell, 300-cycle kit for 2 x 150 bp paired reads). The number of shotgun metagenomic sequences ranged from 55 - 145 million paired reads per sample, with 92% of bases above a quality score of Q30. Adapter sequences and PhiX were removed from all reads with BBDuk (part of the BBTools suite, v35.85 (6)), and quality trimming was performed with fastp (7) using a sliding window of 8 bases and a quality threshold of 28. Reads from each sample were assembled with Megahit (8), and genes were predicted with Prodigal v2.6.3 (9) in meta mode. Megahit was chosen as the assembler for its memory efficiency and speed with complex metagenomes, anticipating that many samples may need to be processed in parallel. Only sample-specific assemblies were performed (i.e. a separate assembly for each sample).

Quality-filtered, unassembled reads were assigned taxonomy with Kaiju and the NCBI nr+euk database (10). Antimicrobial resistance genes (ARGs) were identified among predicted proteins in metagenomic assemblies with AMRFinderPlus v3.12.8 (11). When reporting results, AMRFinder hits labeled as HMM or PARTIALP were excluded, as these are lower confidence results that are often assigned low-specificity annotations such as "bla" (a generic beta lactamase). ARGs were also identified in unassembled short reads (i.e., an assembly-free, read-based approach) with the deeparg short reads pipeline v1.0.4 (12).

The metagenomic sequence coverage of each ARG is reported as the coverage of the assembled contig that encodes it, which is calculated with CoverM (13). Coverages of ARGs are reported at the "gene_symbol" level, as defined by AMRFinderPlus. Coverages are reported in units of transcripts (or fragments) per million (TPM). TPM is a proportional unit (multiplied by one million for convenience) that is normalized to the length of each contig as well as to the total library size. Coverages in units of TPM are considered to be relative, proportional abundances useful for general compositional comparisons among samples. Metagenomic sequence coverages are not absolute abundances, as they report each gene's proportional representation within a sequence library, not its environmental abundance.

All code for the metagenome sequence analysis pipeline, starting with the original sequence reads and ending with plots of sequencing coverage of each identified ARG and MGE, is open source and publicly available (<https://github.com/Brazelton-Lab/UTwastewaterARG>).

To verify the presence of the AR gene targets used by qPCR, 21 isolates were sequenced. Nine isolates were obtained from a wastewater sample collected on January 24, 2023. Twelve additional isolates were recovered and sequenced from a wastewater sample collected on October 17, 2023. Extracted DNA was sequenced at Utah Public Health Laboratory on a MiSeq platform (V2, 300-cycle kit for 2 x 150 bp paired reads). Sequencing yield was 1 - 2 million paired reads per isolate. Quality control, assembly, gene identification, and all analyses with isolate genomes were conducted by Utah Public Health Laboratory with Grandeur v3.0.20230310 (<https://github.com/UPHL-> BioNGS/Grandeur). Grandeur is based on the CDC's PHoeNIx workflow (<https://github.com/CDCgov/phoenix>), with additional quality metrics and gene prediction utilities. Two batches of isolates have been sequenced thus far, one batch of isolates collected during the initial project year and a second batch collected during the first option year. The presence/absence of ARGs in isolates reported here was determined by running AMRFinderPlus on all isolates from both batches with the same version of AMRFinderPlus (v3.12.8).

All data is posted on BioProject accession number PRJNA1280114.

#### Limit of Detection by qPCR and culture based methods

Antimicrobial-resistant bacterial isolates (**Table S1**) from the CDC and FDA Antimicrobial Resistance Isolate Bank were grown in 400 ml LB media with 5% sheep blood and meropenem (either 2 or 8 mg/L). The isolate cultures were shaken at 35°C overnight and centrifuged to pellet the bacteria. The pellet was then resuspended in 2 ml 1X PBS. The suspension was diluted by factors of 10 into 1X PBS and spread on LB plates with meropenem to determine the bacterial concentration. Each concentrated isolate (40 μl) and 500 μl Zymo was added to 44 ml wastewater from student campus residences that include kitchens and common spaces. Ten- and twenty-fold serial dilutions of the suspension were diluted into wastewater (n=5 for each dilution) such that the final volume was 40 ml. Bacteria were precipitated using PEG-6000 and sodium chloride as described above for composite samples. The pellet was resuspended in 1.5 ml PBS and 500 μl was processed for DNA extraction into 50 μl of 10 mM Tris solution for qPCR. At the same time, DNA from 100 μl of concentrated isolate and 5 μl spike-in Zymo was extracted to determine the concentrations of 16S and AR GC in the isolate. DNA in wastewater was extracted using the same method. Concentrations of CP genes and 16S rRNA genes were determined by qPCR using the method described in section “Quantification of genes by qPCR” above. All concentrations were normalized to Zymo recovery.

#### Comparison of sample volumes, handling, and storage

Sampling handling conditions—including hold time, temperature, and processing stage—on gene abundance, was evaluated using a single 4-L grab sample divided into 20 ml to 40 ml aliquots and subjected to five treatment types. First, raw samples were processed immediately to serve as controls (n=3). Second, samples were concentrated using PEG centrifugation (described above), resuspended in 1X PBS, and then held for 1 day at 4°C, ‑20°C or ‑70°C (n=3 per condition). Third, nucleic acids were extracted immediately then stored for 7 days at either -20°C or -70°C (n=3 each), or for 29 days at -20°C (n=6). Fourth, raw aliquots were stored at 4°C, ‑20°C, or -70°C for 1, 7, or 29 days prior to processing, simulating variability in shipping and storage conditions (n=3 per condition; total n=27). Lastly, raw samples were preserved by mixing 1:1 with 50% glycerol in 10 mM 2-amino-2-(hydroxymethyl)propane-1,3-diol (Tris) and stored for 1, 7, or 29 days at 4°C, -20°C, or -70°C before concentration and nucleic acid extraction (n=3 each; total n=27).

The effect of the concentrating 40 or 500-ml of sample on the gene abundance was investigated. Specifically, the treatments included (a) four 40-ml samples centrifuged for 10 minutes at 31,400 *x* *g* and 4°C, (b) two 40-ml samples centrifuged for 1 hour at 3,400 *x* *g* and room temperature, and (c) four 500-ml samples centrifuged at 4,300 *x g* and 4°C for 1 hour. These speeds and temperatures were tested with the understanding that not all laboratories have access to high-speed, temperature-controlled centrifuges. After centrifugation, the samples were processed as described above regarding the nucleic acid extractions.

#### Statistical methods

In the qPCR limit of detection study ANOVA or ANOVA on ranks (when not normally distributed) was used to determine for each trial the mean Ct value for the lowest dilution that was above a Ct of 35 and at a significantly different concentration than the next lowest dilution. The qPCR standard curve calculation was then used for this Ct to estimate the GC/ml. Mean and standard deviation (x̅ ± SD) GC/ml for the three trials was then estimated. The culture detection limit for each of three trials was estimated independently based on linear regressions and a Ct cutoff of 35. Then the average and standard deviation (x̅ ± SD) of these detection limits was estimated. These values are reported in **Table 3**.

Correlations among CP log gene concentrations in composite, passive, and biofilm samples from the rehabilitation hospital after normalization to monthly building water usage rates were determined using Spearman rank order correlations with a significance level of *P* < 0.05. Daily values were calculated for composite samples as log (GC/ml)/(monthly average gal *10^3^/day), for passive samplers as log (GC/g)/(monthly average gal *10^3^/day), and for biofilm samples as log (GC/cm^2^)/(monthly average gal *10^3^/day). Values are reported in **Table 4**, while non normalized values are reported in **Table S5**. Scatterplots of CP gene abundance in each sample type that were found to be significantly correlated by Spearman’s rank order correlations (**Table 4** and **Table S5**) were plotted with SAS (v 9.4) and are shown in **Figure S2** (water usage normalized CP genes) and **Figure S3** (non-normalized log CP genes).

To assess the effects of the independent variables of hold time, hold temperature, hold method, and mixed effects (time and temperature, time and hold method, temperature and hold method, or all three variables) on the dependent variable CP gene and 16S rRNA gene concentrations, we used SAS (v 9.4) general linear mixed effect model. The source of overall model variation was determined as the percent of the total variation of type 1 error for the general linear model for only those independent variables with P < 0.05 for the overall model. These values are reported in **Table 5**.

A nonparametric regression technique was used to smooth the scatter plots of CP genes over time in the wastewater from the rehabilitation hospital as shown in **Figure 3**. The smoothing was conducted in SAS (v 9.4 PROC LOESS) with smoothing parameters from 0.1 to 0.25.

ANOVA was used to determine if there was a difference in the mean log GC/ml of wastewater among 40 ml and 500 ml grab samples as shown in **Figure S1**. This analysis was also conducted for the log CP gene copies/ml of wastewater normalized by the log 16s rRNA gene copies/ml of wastewater to account for any free DNA that was not associated with an intact cell.

To evaluate if passive sampler deployment time influenced the CP gene and 16S rRNA gene abundance, linear regressions of gene abundance versus days deployed were generated in SAS (v 9.4). These plots are shown in **Figure S4**. We also assessed if there were any outliers in the data using a Rosner’s or Dixon’s Outlier Test with a 5% significance level using ProUCL (<https://www.epa.gov/land-research/proucl-software>).

**Supplemental tables**

Table S1. Isolates used to determine limits of detection of qPCR and culture-based methods.

| CDC and FDA AR Isolate Bank number | Organism | Resistance gene | Meropenem resistance (mg/L) |
| --- | --- | --- | --- |
| 0046 | *Klebsiella pneumoniae* | *bla*_VIM-27_  *bla*_OXA-1_ | >8 |
| 0049 | *Klebsiella pneumoniae* | *bla*_NDM-1_  *bla*_OXA-1_ | >8 |
| 0074 | *Klebsiella aerogenes* | *bla*_OXA-48_ | 2 |
| 0090 | *Pseudomonas aeruginosa* | *bla*_KPC-5_  *bla*_OXA-50_ | >8 |
| 0103 | *Pseudomonas aeruginosa* | *bla*_IMP-1_  *bla*_OXA-396_ | >8 |

Table S2. qPCR primer and probe sequences and fluorescent tags.

| Target | Forward (F) primer, Reverse (R) primer, Probe (Pr) (5’-3’) | Reference |
| --- | --- | --- |
| 16S rRNA | F ^b^: TGG AGC ATG TGG TTT AAT TCG A  R ^b^: TGC GGG ACT TAA CCC AAC A Pr ^c^: (Cy3) CA CGA GCT GAC GAC ARC CAT GCA |  |
| *bla*_IMP_ | F ^b^: GGC GG[+A] ^a^AT AGA GTG GCT  R1 ^b^: CCT TAC CGT HTT TTT IAA GMA GBT CAT  R2 ^b^: TTT GTA GCT TGC ACC TTA TTG TCT TT  Pr1 ^c^: (HEX) AY TCT CRR TCH [+A]^a^TY YCM [+A] ^a^ CRT [+A] ^a^T GC  Pr2 ^c^: (CY3) TAT [+G] ^a^C ATC T[+G] ^a^ AAT [+T] ^a^A A[+C] ^a^ AAA T[+G] A | (14) |
| *bla*_KPC_ | F ^b^: GGC CGC CGT GCA ATA C  R ^b^: GCC GCC CAA CTC CTT CA Pr ^c^: (FAM) TG ATA ACG CCG CCG CCA ATT TGT | (3) |
| *bla*_NDM_ | F ^b^: GAC CGC CCA GAT CCT CAA  R ^b^: CGC GAC CGG CAG GTT Pr ^c^: (FAM) TGG ATC AAG CAG GAG AT | (3) |
| *bla*_OXA‑48-like_ | F ^b^: ACG GGC GAA CCA AGC AT R ^b^: GCG ATC AAG CTA TTG GGA ATT T Pr ^c^: (FAM) TTA CCC GCA TCT ACC | (4) |
| *bla*_VIM_ | F ^b^: GAA AAA CAC AGC GGC ACT TCT  R ^b^: CAC GCG TTA CRG GAA GTC CAA Pr ^c^: (HEX) CGG AGA TTG ARA AGC A | (5) |
| Zymo | F ^d^: CCC TGC CTC TTT CCT TAT TGA AG  R ^d^: GCA STA ACA AAC ACC AAC ACT TC Pr ^e^: (HEX) CCC TAG CAA CCC TGG TTG TTA TCG A | Zymo BIOMICs, this work |

^a^ Indicates locked base

^b^ 0.5 µM

^c^ 0.25 µM

^d^ 0.4 µM

^e^ 0.2 µM

Table S3. Positive controls (standards) for qPCR.

| Gene | 5’-3’ sequence for positive control or source of control |
| --- | --- |
| 16S rRNA | Geneblock IDT-DNT  GGT GGA GCA TGT GGT TTA ATT CGA AGC AAC GCG AAG AAC CTT ACC AGG TCT TGA CAT CCC GAT GAC CGC CCT AGA GAT AGG GTT TCT CTT CGG AGC AAC GGT GAC AGG TGG TGC ATG GTT GTC GTC AGC TCG TGT CGT GAG ATG TTG GGT TAA GTC CCG CAA C |
| *bla*_KPC_ | Geneblock IDT-DNT  TGA CAA CAG GCA TGA CGG TGG CGG AGC TGT CCG CGG CCG CCG TGC AAT ACA GTG ATA ACG CCG CCG CCA ATT TGT TGC TGA AGG AGT TGG GCG GCC CGG CCG GGC TGA CGG CCT TCA TGC GCT CTA TCG GCG ATA CCA CG |
| *bla*_NDM_ | Geneblock IDT-DNT  CCG ATG ACC AGA CCG CCC AGA TCC TCA ACT GGA TCA AGC AGG AGA TCA ACC TGC CGG TCG CGC TGG CGG TGG TGA CTC ACG CGC ATC AGG ACA AGA TGG GCG GTA TGG ACG CGC TGC ATG CGG CG |
| *bla*_VIM_ | Geneblock IDT-DNT  GAA AAA CAC AGC GGC ACT TCT TTT TTC GGA GAT TGA RAA GCA TTT TTT TGG ACT TCC YGT AAC GCG TG |
| *bla*_OXA-48-like_ | Geneblock IDT-DNT  GAA CAT AAA TCA CAG GGC GTA GTT GTG CTC TGG AAT GAG AAT AAG CAG CAA GGA TTT ACC AAT AAT CTT AAA CGG GCG AAC CAA GCA TTT TTA CCC GCA TCT ACC TTT AAA ATT CCC AAT AGC TTG ATC GCC CTC GAT TT |
| *bla*_IMP_ | Extraction of DNA from *Klebsiella pneumoniae* culture derived from CDC and FDA AR Isolate Bank, AR0034, accension number SAMN04014875. |
| Zymo | Geneblock IDT-DNT  ATC CCT GCC TCT TTC CTT ATT GAA GTT AAT ACC CAA AAC ACT TAA ACA AAA ATA ACC AGT TGT TAT AAG AAC ATT TGT TCC TTG TTA GTT GAA TTA ATG CTA TAA TAT TTA TAC CCC CTA GCA ACC CTG GTT GTT ATC GAG AAG AAG ATT CTT TCT TCT GGA AAG GGG GTG TGT AGA TAT GGA AGT GTT GGT GTT TGT TAC TGC GG |

Table S4. qPCR reaction efficiencies for single-plex and duplex assays and number (n) of log dilutions in the standard curve. 16S is 16S rRNA; KPC is *bla*_KPC_; NDM is *bla*_NDM_ VIM is *bla*_VIM_; OXA is *bla*_OXA-48-like_; and IMP is *bla*_IMP_.

| Single-plex (fluor) | Efficiency (%) | n |  | Duplex (fluor) | Efficiency (%) | n |
| --- | --- | --- | --- | --- | --- | --- |
| Zymo (HEX) | 96.0 | 7 |  | Zymo (HEX) with *C. auris* | 88.9 | 7 |
| KPC (FAM) | 94.0 | 7 |  | KPC (FAM) with 16S | 92.6 | 7 |
| 16S (Cy3) | 99.1 | 7 |  | 16S (CY3) with KPC | 119.0 | 7 |
| NDM (VIC) | 115.6 | 5 |  | NDM (VIC) with OXA | 97.3 | 7 |
| NDM (FAM) | 98.7 | 7 |  | NDM (FAM) with IMP | 110.5 | 6 |
| IMP (CY3) | 93.1 | 6 |  | IMP (CY3) with NDM | 96.3 | 6 |
| VIM (HEX) | 90.5 | 8 |  | VIM (HEX) with OXA | 96.7 | 7 |
| OXA (FAM) | 100.1 | 7 |  | OXA (FAM) with NDM OXA (FAM) with VIM | 97.8 97.8 | 7 7 |

Table S5. Spearman rank order correlations among *bla*_KPC_, *bla*_OXA‑48-like_, and *bla*_VIM_, log_10_ gene concentrations for composite, passive samplers, and sewer biofilm sample collection methods.^a,b^

|  |  | Composite (C) | | | Passive (P) | | | Sewer biofilm (B) | | |
| --- | --- | --- | --- | --- | --- | --- | --- | --- | --- | --- |
|  |  | *bla*_KPC_ | *bla*_OXA_ | *bla*_VIM_ | *bla*_KPC_ | *bla_OXA-48-like_* | *bla*_VIM_ | *bla*_KPC_ | *bla*_OXA-48-like_ | *bla_VIM_* |
| C | *bla*_KPC_ | -- | 0.63  <0.001  [59] ^b^ | 0.56  0.007  [22] | NS ^c^ | NS ^d^ | 0.65  0.008  [15] | NS | NS | NS |
|  | *bla*_OXA-48-like_ |  | -- | 0.60  0.003  [22] | NS | NS | 0.65  0.009  [15] | NS | NS | NS |
|  | *bla*_VIM_ |  |  | -- | NS | NS | 0.78  0.005  [10] | NS | NS | NS |
| P | *bla*_KPC_ |  |  |  | -- | 0.42  0.002  [52] | NS | NS | NS | NS |
|  | *bla*_OXA-48-like_ |  |  |  |  | -- | NS | NS | NS | NS |
|  | *bla*_VIM_ |  |  |  |  |  | -- | -0.8  0.01  [7] | NS | NS |
| B | *bla*_KPC_ |  |  |  |  |  |  | -- | 0.76  <0.001  [15] | NS |
|  | *bla*_OXA-48-like_ |  |  |  |  |  |  |  | -- | NS |
|  | *bla*_VIM_ |  |  |  |  |  |  |  |  | -- |

^a.^ *bla*_NDM_ and *bla*_IMP_ were excluded as they were not correlated or correlated in ≤ 5 fewer samples.

^b^ Spearman’s rho, *P*, [number of samples]

^c^ NS = not significant at a *P* < 0.05 level

Table S6. Bacterial concentrations [log10(colony forming units/100 ml)] determined by CHROMagar KPC and MacConkey I. ^a^

|  | CHROMagar KPC | | | MacConkey I | | |
| --- | --- | --- | --- | --- | --- | --- |
| Date | E. coli | Klebsiella, Enterobacter, Citrobacter | Pseudomonas, Acinetobacter | | Lactose fermenting gram negative | Non-lactose fermenters |
| Composite samples | | | | | | |
| 1/24/2023 | 6.57 | 8.35 | 7.02 | | 8.24 | 8.47 |
| 2/9/2023 | 7.00 | 7.66 | 8.18 | | NA | 8.70 |
| 3/22/2023 | NA | NA | NA | | 9.52 | 8.54 |
| 3/29/2023 | NA | NA | NA | | 9.81 | 9.76 |
| 4/5/2023 | 7.90 | 9.14 | 8.28 | | 9.19 | 9.54 |
| 4/19/2023 | 8.82 | 9.09 | 8.32 | | 9.56 | 9.26 |
| 4/26/2023 | 9.49 | 10.06 | 8.95 | | 11.06 | 10.48 |
| 5/22/2023 | 9.86 | 9.60 | 8.78 | | 10.79 | 10.64 |
| 6/21/2023 | 9.30 | 9.90 | 9.60 | | 10.76 | 9.78 |
| 7/17/2023 | 9.34 | 8.78 | 9.08 | | 10.80 | 9.90 |
| 8/14/2023 | NA | 8.00 | 8.00 | | 9.49 | 8.78 |
| 9/19/2023 | 8.08 | 8.72 | 8.32 | | 9.87 | 8.60 |
| 10/17/2023 | 8.41 | 8.76 | 8.41 | | 9.03 | 8.49 |
| 11/15/2023 | 8.49 | 8.69 | 7.90 | | 9.58 | 8.85 |
| x ± SD | 8.48 ± 1.04 | 8.90 ± 0.72 | 8.40 ± 0.66 | | 9.82 ± 0.82 | 9.27 ± 0.75 |
| Moore samples | | | | | | |
| 2/9/2023 | 7.51 | 8.92 | 6.78 | | 8.57 | 8.43 |
| 4/26/2023 | 10.26 | 10.75 | NA | | 10.30 | NA |
| 5/22/2023 | 10.92 | 10.76 | 9.30 | | 11.52 | 10.78 |
| 6/21/2023 | 10.04 | 9.30 | 9.30 | | 10.92 | 9.60 |
| 7/17/2023 | 9.30 | 10.23 | 9.30 | | 11.04 | 10.18 |
| 8/14/2023 | 9.36 | 8.90 | 8.00 | | 10.29 | 8.85 |
| 9/19/2023 | 7.76 | 8.17 | 8.04 | | 8.97 | 7.48 |
| 10/17/2023 | 9.20 | 9.80 | 8.36 | | 9.88 | 10.08 |
| 11/15/2023 | 9.95 | 9.85 | 8.00 | | 10.45 | 9.48 |
| x ± SD | 9.37 ± 1.12 | 9.63 ± 0.88 | 8.39 ± 0.89 | | 10.22 ± 0.96 | 9.36 ± 1.06 |
| Grab sample | | | | | | |
| 2/9/2023 | 6.00 | 6.85 | 7.36 | | 6.78 | 8.15 |
| Biofilm sample | | | | | | |
| 2/9/2023 | 5.90 | 6.30 | 6.85 | | 7.48 | 7.60 |

^a^ Culture colors on CHROMagar KPC agar are assigned as: pink is presumed to be *E. coli*; blue is presumed to be *Klebsiella, Enterobacter* or *Citrobacter*; and white/clear/translucent is presumed to be *Acinetobacter* or *Pseudomonas*. On MacConkey I agar, lactose-fermenting, gram-negative bacterial colonies are red whereas the non-fermenting colonies are white.

Table S7. Pearson correlations among culture log_10_ CFU/ml from CHROMagar KPC and MacConkey I for autosampler composite and passive samplers.

|  |  | Composite | | | | |
| --- | --- | --- | --- | --- | --- | --- |
|  |  | *E. coli* | *Klebsiella, Enterobacter,* and *Citrobacter* | *Pseudomonas* and *Acinetobacter* | Lactose fermenting gram negative | Non-lactose fermenters |
| Passive sampler | *E. coli* | 0.89,  0.004  [8] | 0.73,  0.03  [9] | NS ^b^ | NS | 0.73,  0.03  [9] |
|  | *Klebsiella, Enterobacter,* and *Citrobacter* | -- | NS | NS | NS | 0.88,  0.01  [9] |
|  | *Pseudomonas* and *Acinetobacter* | -- | -- | 0.77,  0.02  [8] | 0.87,  0.01  [7] | 0.78,  0.02  [8] |
|  | Lactose fermenting gram negative | -- | -- | -- | NS | 0.721,  0.03  [9] |
|  | Non-lactose fermenters | -- | -- | -- | -- | NS |

^a^ Pearson’s r, *P*, [number of samples]

^b^ NS = not significant at a *P* < 0.05 level

Table S8. Contingency tables for detection of carbapenem-resistant CP genes by qPCR in samples collected by autosampler composite, passive sampler, and sewer biofilm.

| *bla*_VIM_ | | | | | | | | | | | |
| --- | --- | --- | --- | --- | --- | --- | --- | --- | --- | --- | --- |
|  |  | Composite | |  |  | Biofilm | |  |  | Composite | |
|  |  | Pos | Neg |  |  | Pos | Neg |  |  | Pos | Neg |
| Passive | Pos | 10 [18.9%] | 5  [9.4%] | Passive | Pos | 3  [20%] | 4 [26.7%] | Biofilm | Pos | 2 [12.5%] | 1 [6.3%] |
|  | Neg | 11 [20.8%] | 27 [50.9%] |  | Neg | 0  [0%] | 8 [53.3%] |  | Neg | 4  [25%] | 9 [56.3%] |
| *bla*_OXA-48-like_ | | | | | | | | | | | |
|  |  | Composite | |  |  | Biofilm | |  |  | Composite | |
|  |  | Pos | Neg |  |  | Pos | Neg |  |  | Pos | Neg |
| Passive | Pos | 51 [96.2%] | 1  [1.9%] | Passive | Pos | 15 [100%] | 0  [0%] | Biofilm | Pos | 16 [100%] | 0  [0%] |
|  | Neg | 1  [1.9%] | 0  [0%] |  | Neg | 0  [0%] | 0  [0%] |  | Neg | 0  [0%] | 0  [0%] |
| *bla*_NDM_ | | | | | | | | | | | |
|  |  | Composite | |  |  | Biofilm | |  |  | Composite | |
|  |  | Pos | Neg |  |  | Pos | Neg |  |  | Pos | Neg |
| Passive | Pos | 3  [5.7%] | 1  [1.9%] | Passive | Pos | 1 [6.7%] | 2 [13.3%] | Biofilm | Pos | 1  [6.3%] | 1  [6.3%] |
|  | Neg | 1  [1.9%] | 48 [90.6%] |  | Neg | 1 [6.7%] | 11 [73.3%] |  | Neg | 2 [12.5%] | 12 [75%] |
| *bla*_KPC_ | | | | | | | | | | | |
|  |  | Composite | |  |  | Biofilm | |  |  | Composite | |
|  |  | Pos | Neg |  |  | Pos | Neg |  |  | Pos | Neg |
| Passive | Pos | 53 [100%] | 0  [0%] | Passive | Pos | 15 [100%] | 0  [0%] | Biofilm | Pos | 16 [100%] | 0  [0%] |
|  | Neg | 0  [0%] | 0  [0%] |  | Neg | 0  [0%] | 0  [0%] |  | Neg | 0  [0%] | 0  [0%] |
| *bla*_IMP_ | | | | | | | | | | | |
|  |  | Composite | |  |  | Biofilm | |  |  | Composite | |
|  |  | Pos | Neg |  |  | Pos | Neg |  |  | Pos | Neg |
| Passive | Pos | 1  [1.9%] | 2  [3.8%] | Passive | Pos | 1 [6.7%] | 1  [6.7%] | Biofilm | Pos | 1  [6.7%] | 0  [0%] |
|  | Neg | 0  [0%] | 50 [94.3%] |  | Neg | 0  [0%] | 13 [86.7%] |  | Neg | 0  [0%] | 15 [93.8%] |

Table S9. Diversity indices for metagenomes from autosampler composite, passive samples, sewer biofilm, and grab samples.

| Sample type ^a^ | Number of ARG taxa or richness, S | Dominance, D | Simpsons, 1-D | Shannon, H | Evenness, E |
| --- | --- | --- | --- | --- | --- |
| Composite 1, 2023 | 212 | 0.035 | 0.97 | 4.1 | 0.28 |
| Composite 2, 2023 | 165 | 0.035 | 0.97 | 4.1 | 0.36 |
| Composite 3, 2024 | 226 | 0.020 | 0.98 | 4.6 | 0.44 |
| Grab, 2023 | 155 | 0.06 | 0.94 | 3.6 | 0.25 |
| Passive 1, 2023 | 218 | 0.03 | 0.97 | 4.2 | 0.29 |
| Passive 2, 2024 | 203 | 0.03 | 0.97 | 4.1 | 0.31 |
| Biofilm, 2023 | 155 | 0.05 | 0.95 | 3.8 | 0.27 |

^a^ All samples in either 2023 or 2024 were collected on the same day.

Table S10. Contingency tables qPCR versus Cepheid GeneXpert Carba-R assay for autosampler collected samples.

| *bla*_KPC_ | qPCR positive | qPCR negative | Sum of samples |
| --- | --- | --- | --- |
| Carba-R positive | 60 [100%] | 0 [0%] | 60 |
| Carba-R negative | 0 [0%] | 0 [0%] | 0 |
| Sum of samples | 60 | 0 | 60 |
| *bla*_VIM_ |  |  |  |
| Carba-R positive | 22 [36.7%] | 38 [63.3%] | 60 |
| Carba-R negative | 0 [0%] | 0 [0%] | 0 |
| Sum of samples | 22 | 38 | 60 |
| *bla*_OXA-48-like_ |  |  |  |
| Carba-R positive | 59 [98.3%] | 1 [1.7%] | 60 |
| Carba-R negative | 0 [0%] | 0 [0%] | 0 |
| Sum of samples | 59 | 1 | 60 |
| *bla*_NDM_ |  |  |  |
| Carba-R positive | 2 [3.3%] | 0 [0%] | 2 |
| Carba-R negative | 2 [3.3%] | 56 [93.3%] | 58 |
| Sum of samples | 4 | 56 | 60 |
| *bla*_IMP_ |  |  |  |
| Carba-R positive | 0 [0%] | 0 [0%] | 0 |
| Carba-R negative | 1 [1.7%] | 59 [93.3%] | 60 |
| Sum of samples | 1 | 59 | 60 |

Table S11. Contingency tables qPCR versus Cepheid GeneXpert Carba-R assay for passive sampler collected samples

| *bla*_KPC_ | qPCR positive | qPCR negative | Sum of samples |
| --- | --- | --- | --- |
| Carba-R positive | 53 [100%] | 0 [0%] | 53 |
| Carba-R negative | 0 [0%] | 0 [0%] | 0 |
| Sum of samples | 53 | 0 | 53 |
| *bla*_VIM_ |  |  |  |
| Carba-R positive | 15 [28.3%] | 38 [71.7%] | 53 |
| Carba-R negative | 0 [0%] | 0 [0%] | 0 |
| Sum of samples | 15 | 38 | 53 |
| *bla*_OXA-48-like_ |  |  |  |
| Carba-R positive | 52 [98%] | 1 [1.9%] | 53 |
| Carba-R negative | 0 [0%] | 0 [0%] | 0 |
| Sum of samples | 52 | 1 | 53 |
| *bla*_NDM_ |  |  |  |
| Carba-R positive | 2 [3.8%] | 2 [3.8%] | 4 |
| Carba-R negative | 2 [3.8%] | 47 [88.7%] | 49 |
| Sum of samples | 4 | 49 | 53 |
| *bla*_IMP_ |  |  |  |
| Carba-R positive | 0 [0%] | 0 [0%] | 0 |
| Carba-R negative | 3 [56.6%] | 50 [94.3%] | 53 |
| Sum of samples | 3 | 50 | 53 |

Table S12. Contingency tables qPCR versus Cepheid GeneXpert Carba-R assay for sewer biofilm samples

| *bla*_KPC_ | qPCR positive | qPCR negative | Sum of samples |
| --- | --- | --- | --- |
| Carba-R positive | 16 [100%] | 0 [0%] | 16 |
| Carba-R negative | 0 [0%] | 0 [0%] | 0 |
| Sum of samples | 16 | 0 | 16 |
| *bla*_VIM_ |  |  |  |
| Carba-R positive | 3 [18.8%] | 0 [0%] | 3 |
| Carba-R negative | 0 [0%] | 13 [81.2%] | 13 |
| Sum of samples | 3 | 13 | 16 |
| *bla*_OXA-48-like_ |  |  |  |
| Carba-R positive | 16 [100%] | 0 [0%] | 16 |
| Carba-R negative | 0 [0%] | 0 [0%] | 0 |
| Sum of samples | 16 | 0 | 16 |
| *bla*_NDM_ |  |  |  |
| Carba-R positive | 0 [0%] | 3 [18.8%] | 3 |
| Carba-R negative | 2 [12.5%] | 11 [68.8%] | 13 |
| Sum of samples | 2 | 14 | 16 |
| *bla*_IMP_ |  |  |  |
| Carba-R positive | 0 [0%] | 0 [0%] | 0 |
| Carba-R negative | 1 [6.3%] | 15 [93.8%] | 16 |
| Sum of samples | 1 | 15 | 16 |

Table S13. Antibiotic use in the facility (antimicrobial days per 1000 days present) during the 16 month observation period. Imipemen was not perscribed during the observation period.

| Month-Year | Ertapenem | Meropenem |
| --- | --- | --- |
| 01-2023 | 0 | 5.7 |
| 02-2023 | 0 | 28 |
| 03-2023 | 0 | 20.3 |
| 04-2023 | 0 | 18.8 |
| 05-2023 | 5.04 | 18.1 |
| 06-2023 | 0 | 18.5 |
| 07-2023 | 8.1 | 40.4 |
| 08-2023 | 8.1 | 2.8 |
| 09-2023 | 2.7 | 17.7 |
| 10-2023 | 0 | 2.8 |
| 11-2023 | 0 | 0 |
| 12-2023 | 0 | 50.1 |
| 01-2024 | 0 | 136.5 |
| 02-2024 | 0 | 0 |
| 03-2024 | 0 | 0 |

Table S14. Percentage of patients in the facility who were toileting or showering.

| Month-Year | Patient days | Percentage toileting | Percentage showering |
| --- | --- | --- | --- |
| 01-2023 | 1764 | unknown | unknown |
| 02-2023 | 1804 | 37% | 16% |
| 03-2023 | 1917 | 30% | 14% |
| 04-2023 | 1377 | 25% | 13% |
| 05-2023 | 1482 | 38% | 17% |
| 06-2023 | 1338 | 38% | 8% |
| 07-2023 | 1423 | 36% | 19% |
| 08-2023 | 1440 | 30% | 13% |
| 09-2023 | 1402 | 36% | 19% |
| 10-2023 | 1373 | 35% | 13% |
| 11-2023 | 1299 | 38% | 14% |
| 12-2023 | 1234 | 33% | 14% |
| 01-2024 | 1356 | 53% | 17% |
| 02-2024 | 1231 | 45% | 19% |
| 03-2024 | 1498 | 33% | 19% |
| 04-2024 | 1377 | 35% | 13% |

**Supplemental figures**

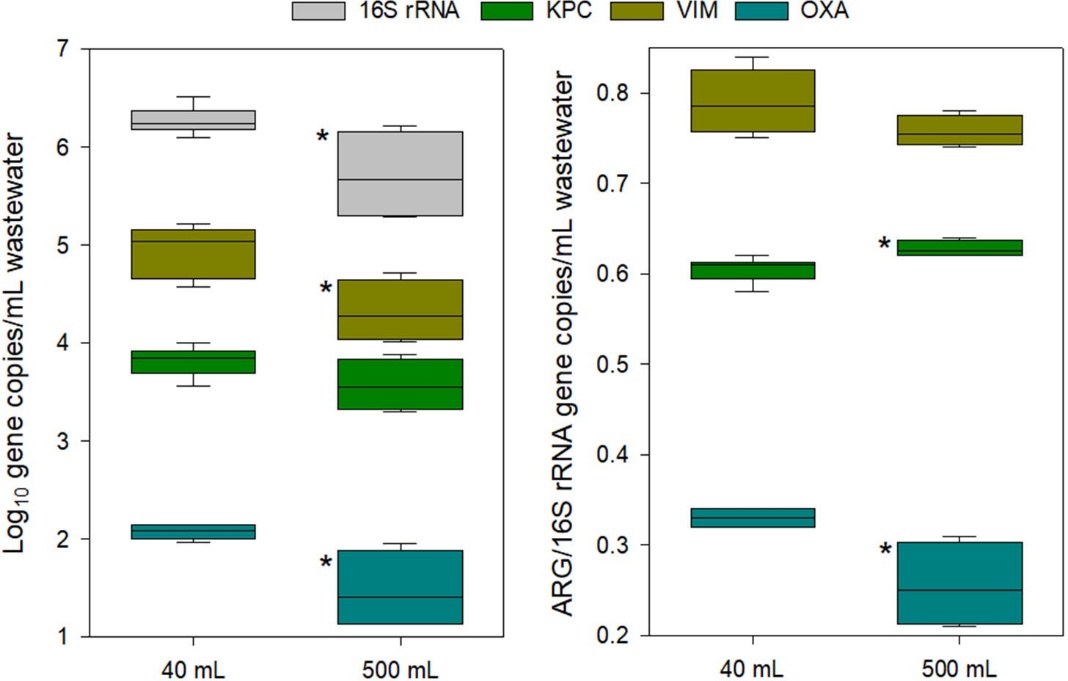

Figure S1. Log gene copies/ml of wastewater when 40 ml (n = 6) or 500 ml (n = 4) of grab sample was processed. Left panel is the absolute abundance observed and right panel is the antimicrobial resistant genes (ARG)/16S rRNA gene abundance. Mean log gene abundance among similar gene targets that were significantly different are labeled with a star. Box plots indicate the first and third quartiles with the whiskers indicating the minimum and maximum values, while the median is shown within the box. Outliers were not present.

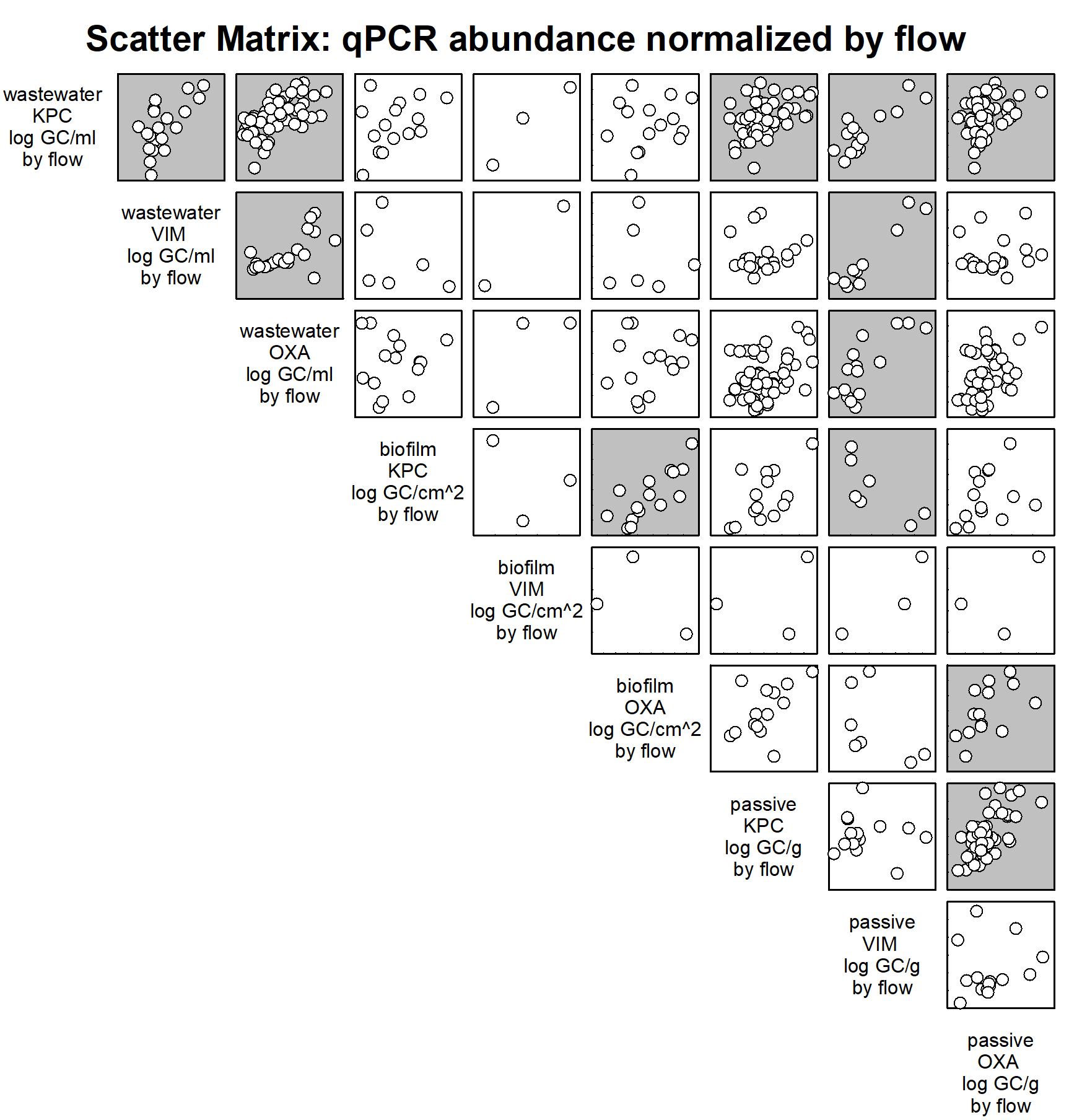

Figure S2. qPCR abundance of *bla*_KPC_, *bla*_OXA‑48-like_, and *bla*_VIM_ genes found to be significantly correlated in autosampler, passive sampler, or sewer biofilms normalized by water usage or flow. Greyed out boxes correspond to signficant correlations based on Spearman’s rank order correlations in Table 4.

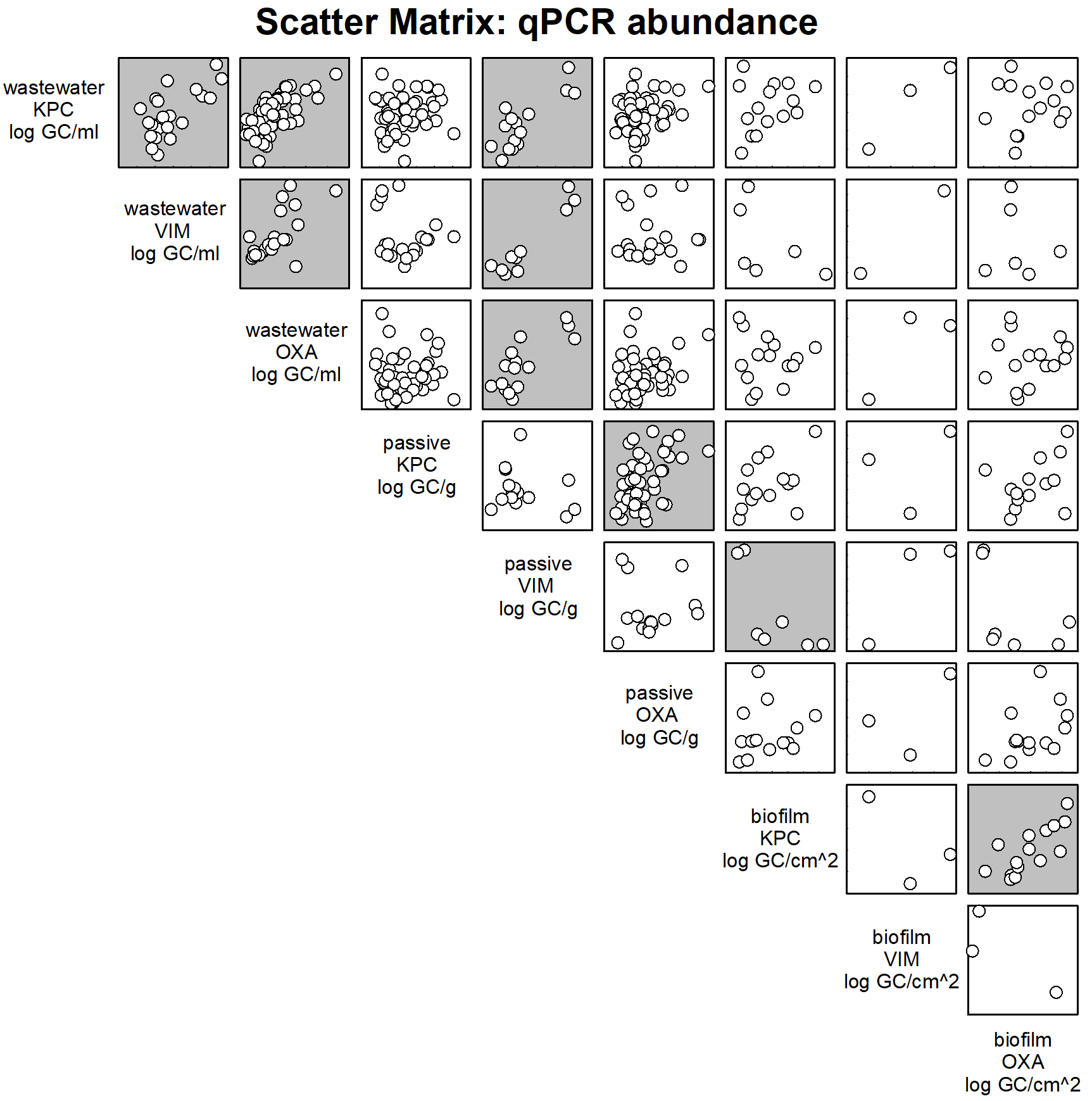

Figure S3. qPCR abundance of *bla*_KPC_, *bla*_OXA‑48-like_, and *bla*_VIM_ genes found to be significantly correlated in autosampler, passive sampler, or sewer biofilms. Greyed out boxes correspond to signficant correlations based on Spearman’s rank order correlations in Table S5.

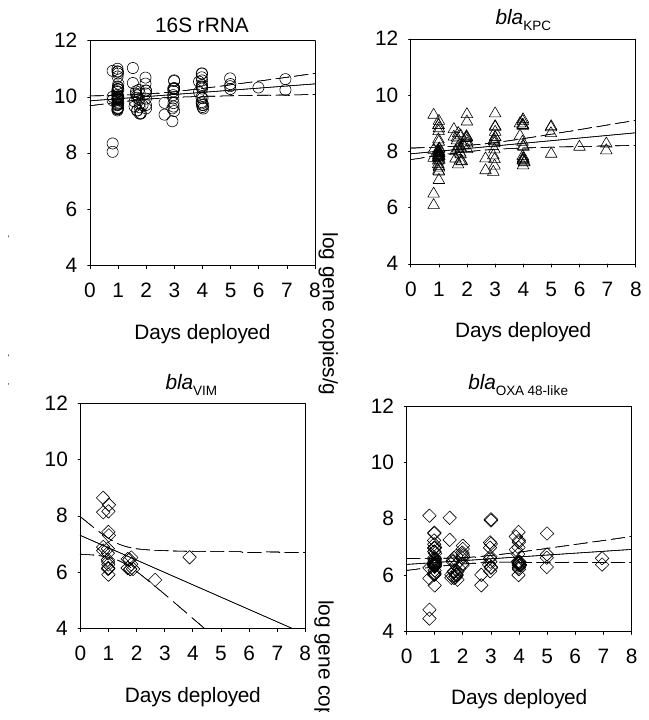

Figure S4. Log gene copies per g of passive sampler versus days the passive sampler was deployed in the sewer system for the duration of the study. Dashed lines indicate the 95% confidence limits on the linear regressions. All R^2 are less 0.15.

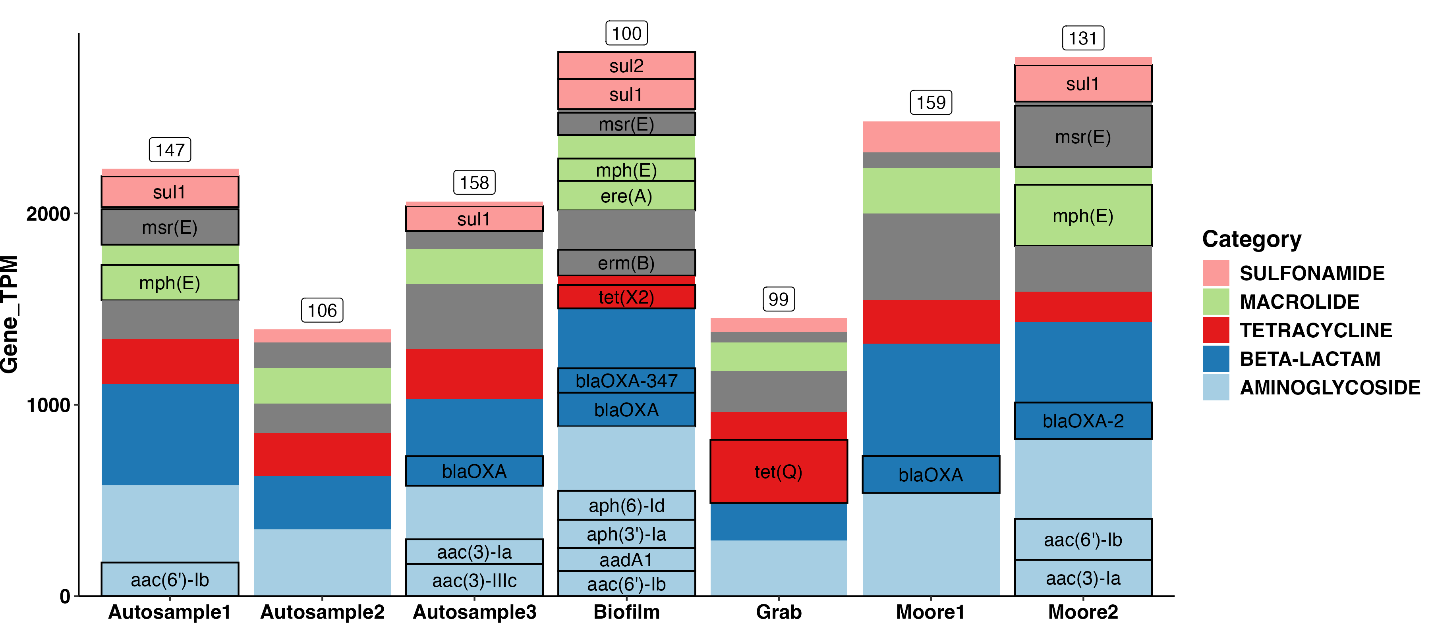

Figure S5. ARGs identified in seven metagenomes, classified according to their categories defined by AMRFinder results labeled as HMM or PARTIALP. Sequencing coverages are reported as TPM (transcripts/fragments per million), a proportional unit normalized to contig length and metagenomic library size. Individual genes are labeled where the size of the bar section is large enough to accommodate text. Unlabeled regions indicate many different genes at low sequence coverage. The total number of ARGs identified in each metagenome is reported in the text box at the top of each column.

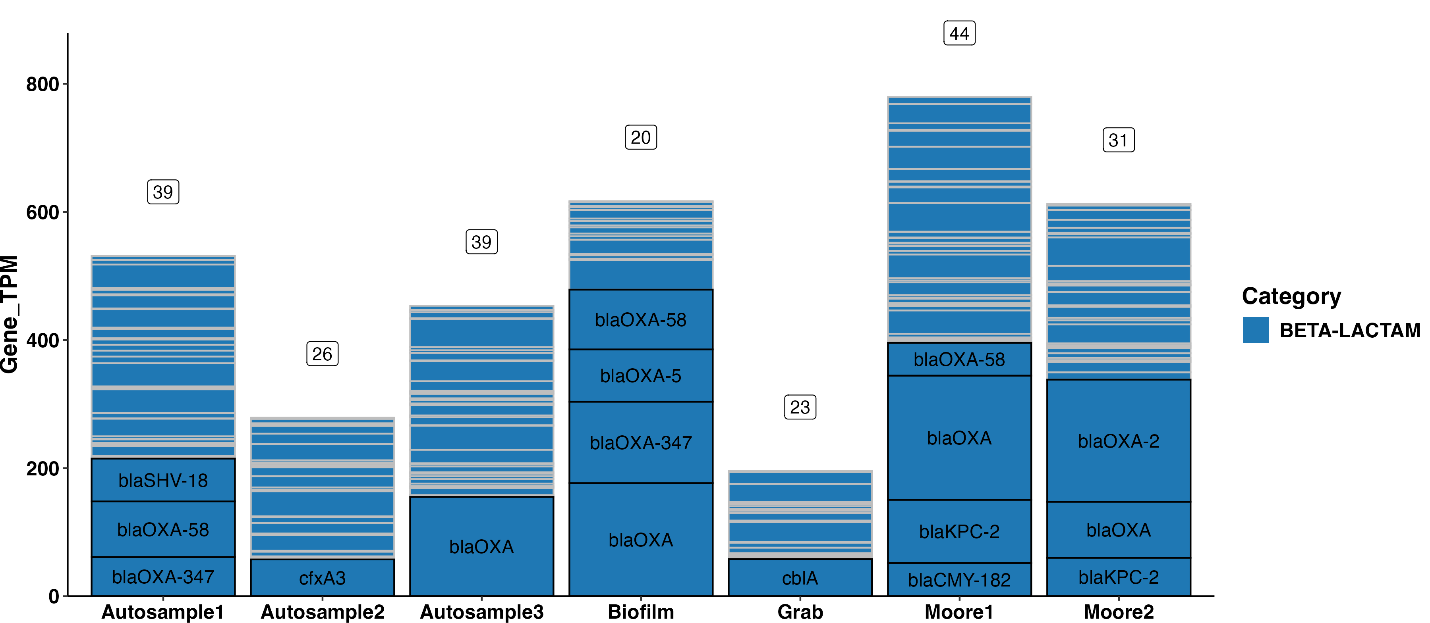

Figure S6. Beta-lactamase genes, excluding AMRFinderPlus results labeled as HMM or PARTIALP. Unlabeled regions indicate many different genes at low sequence coverage. The number of distinct genes identified as "Beta-lactam" by AMRFinder is reported in the text box at the top of each column. All bla genes of interest were detected in at least one sample type, but may be at very low TPM and not identifiable on this figure scale.

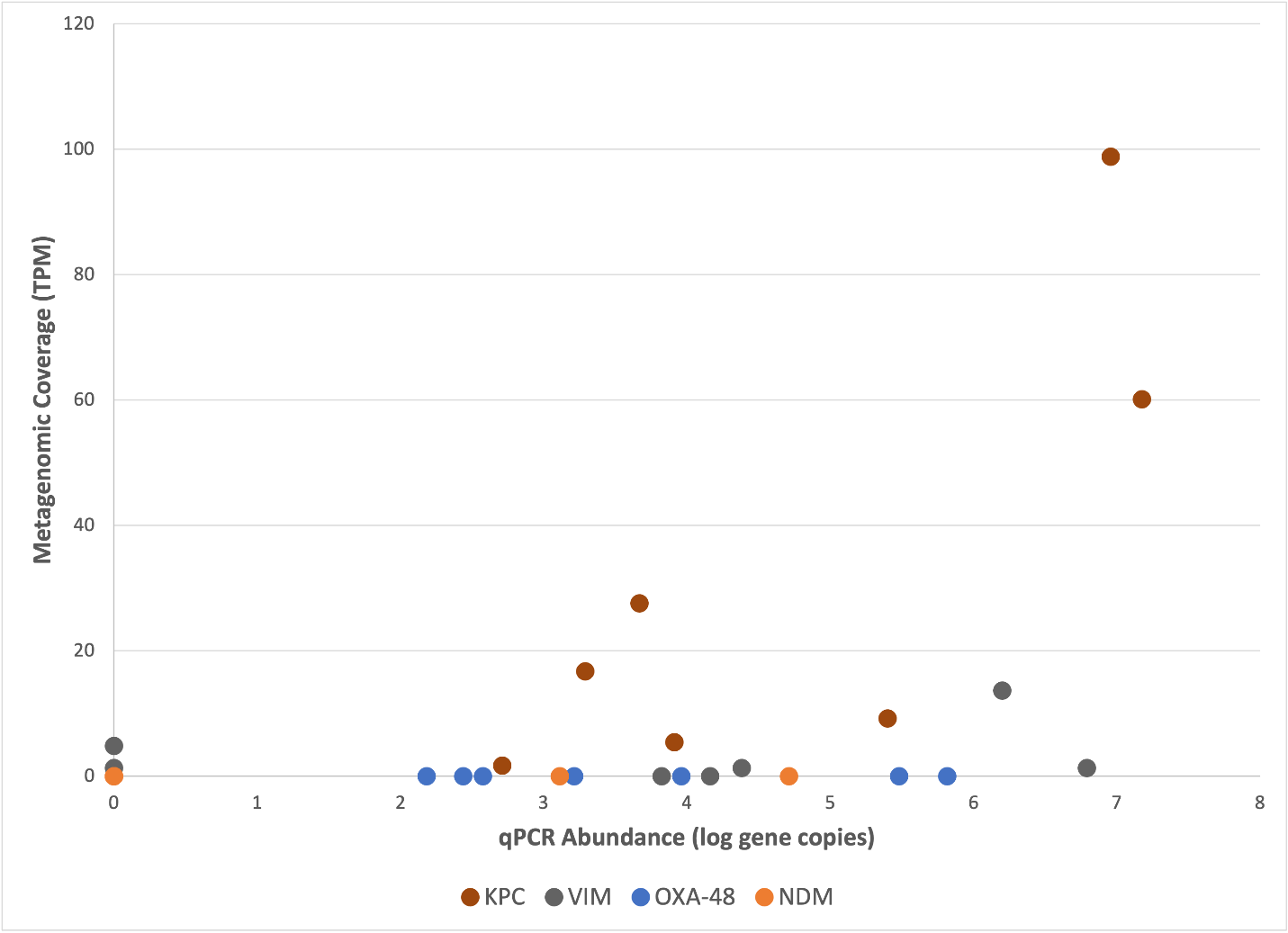

Figure S7. A comparison of metagenomic sequence coverage (cumulative transcripts (or fragments) per million (TPM)) and qPCR abundance (log gene copies) for bla_KPC_, bla_VIM_, bla_OXA‑48-like_, and bla_NDM_. The two methods are significantly correlated (R^2^ = 0.67) for bla_KPC_. However, bla_OXA-48-like_ and bla_NDM_ genes were not detected in any metagenomes.

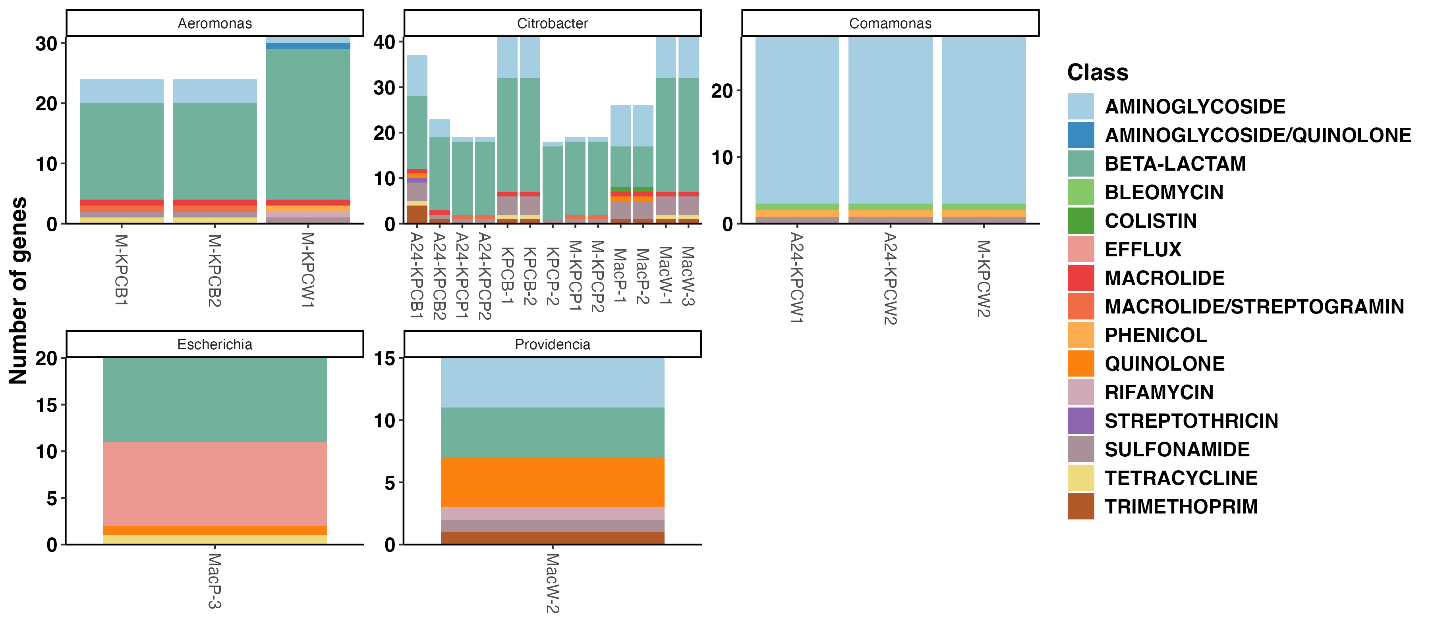

Figure S8. Number of antimicrobial resistance genes (ARGs) identified in the 21 isolates recovered to date. Genes are color-coded according to their class as defined by AMRFinder.

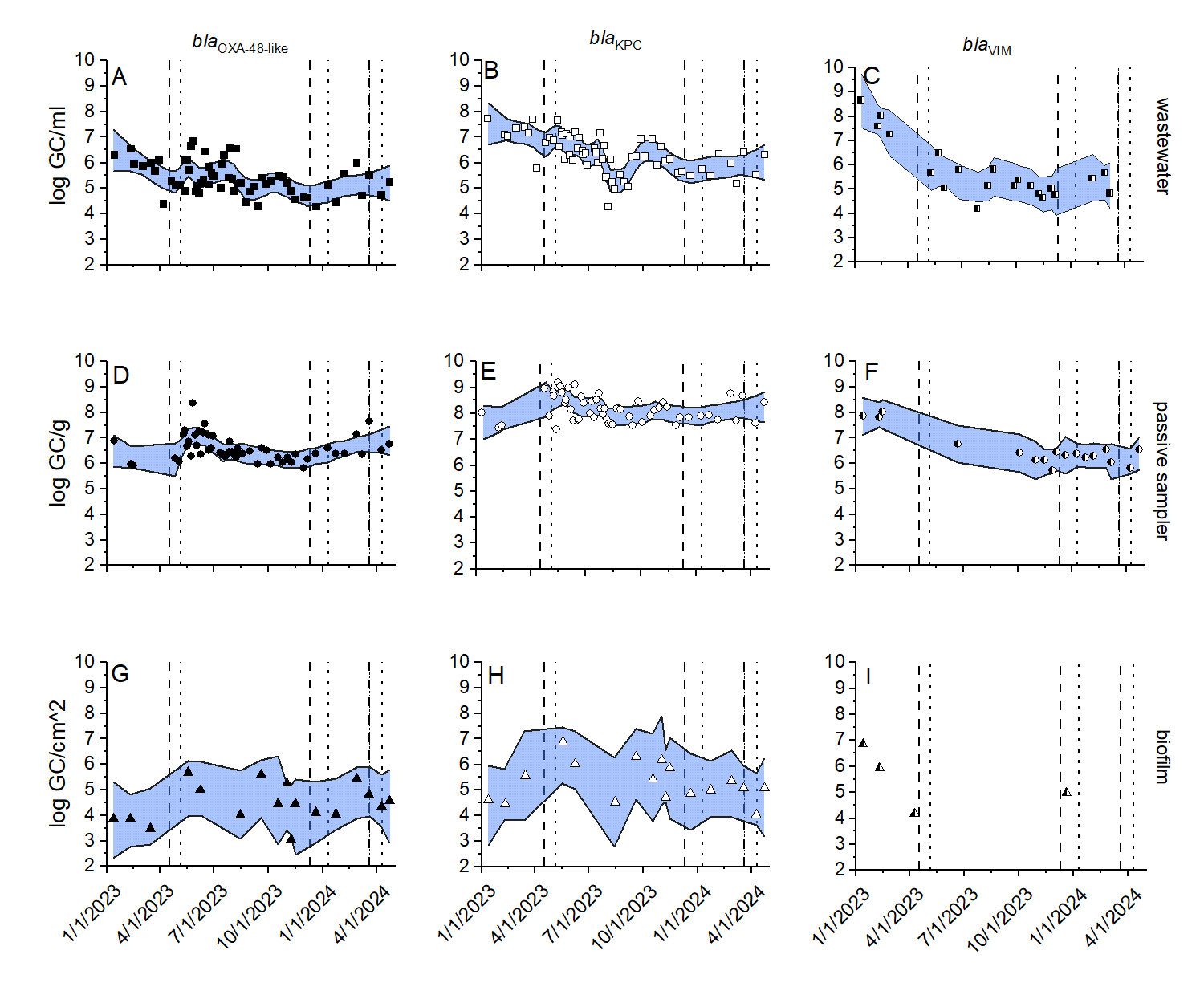

Figure S9. Gene copies (GCs) of *bla*_OXA‑48-like_ (left column, closed symbols), *bla*_KPC_ (center column, open symbols), and *bla*_VIM_ (right column, half-filled symbols) in wastewater collected by autosamplers (top row, circles), passive samplers (middle row, squares), or swabs of sewer biofilm (bottom row, triangles), from January 2023 to April 2024. Non parametric regression smoothing factors range from 0.1 to 0.25. Dash and dotted vertical lines indicate periods when patients infected with CRE or CRPA were in the facility including *Klebsiella aerogenes*, *Enterobacter cloacae* and *Pseudomonas aeruginosa* in period 1 (April, 2023), *P. aerugniosa* in period 2 (December 2023) and *Escherichia coli* and *Citrobacter freundii* in period 3 (March 2024). The CRE infections in April 2023 were not CPO based on PCR. The CRE infection in March 2024 was not tested for CP genes by PCR. None of the CRPA were tested for CP genes.

14.	UPHL. Multiplex Real-Time PCR for the Detection of IMP Genes: Utah Public Health Laboratory; 2022. Report No.: 110395.86.
